# The transcription factor Zt107320 affects the dimorphic switch, growth and virulence of the fungal wheat pathogen *Zymoseptoria tritici*

**DOI:** 10.1101/595454

**Authors:** Michael Habig, Sharon Marie Bahena-Garrido, Friederike Barkmann, Janine Haueisen, Eva Holtgrewe Stukenbrock

## Abstract

*Zymoseptoria tritici* is a filamentous fungus causing Septoria tritici blotch in wheat. The pathogen has a narrow host range and infections of grasses other than susceptible wheat are blocked early after stomatal penetration. During these abortive infections the fungus shows a markedly different expression pattern. However, the underlying mechanisms causing differential gene expression during host and non-host interaction are largely unknown, but likely include transcriptional regulators responsible for the onset of an infection program in compatible hosts. In the rice blast pathogen *Magnaporthe oryzae*, MoCOD1, a member of the fungal Zn(II)_2_Cys_6_ transcription factor family, has been shown to directly affect pathogenicity. Here, we analyse the role of the putative transcription factor Zt107320, a homolog of MoCOD1, during infection of compatible and incompatible hosts by *Z. tritici*. We show for the first time that *Zt107320* is differentially expressed in host versus non-host infections and that lower expression corresponds to an incompatible infection of non-hosts. Applying reverse genetics approaches we further show that Zt107320 regulates the dimorphic switch as well as the growth rate of *Z. tritici* and affects fungal cell wall composition *in vitro*. Moreover, Δ*Zt107320* mutants showed reduced virulence during compatible infections of wheat. We conclude that Zt107320 directly influences pathogen fitness and propose that Zt107320 regulates growth processes and pathogenicity during infection. Our results suggest that this putative transcription factor is involved in discriminating compatible and non-compatible infections.

## Introduction

The fungus *Zymoseptoria tritici* (synonym *Mycosphaerella graminicola*) infects wheat and causes the disease Septoria tritici blotch. The pathogen is found worldwide where wheat is grown and can cause severe reduction in yield (Fones and Gurr, 2015). Upon infection, the fungus enters the leaf through stomata and establishes a hyphal network in the mesophyll. It propagates without causing visual symptoms for 7-14 days before inducing necrosis and producing pycnidia – asexual fructifications, where pycnidiospores are produced that can spread via contact or rain-splash to neighbouring leaves (Brading *et al.* 2002; Kema, G. H. J. *et al.* 1996; Ponomarenko A. S.B. Goodwin, G.H.J. Kema, 2011). *Z. tritici* has a heterothallic mating system and meiosis leads to the production of wind-borne ascospores that are considered to be the main primary inoculum (Kema *et al.* 1996; Morais *et al.* 2016; Ponomarenko A. S.B. Goodwin, G.H.J. Kema, 2011). Recently, we could show that chromosomal inheritance is characterised by frequent chromosome losses and rearrangements during mitosis and a drive of accessory chromosomes during meiosis (Habig *et al.* 2018; Möller *et al.* 2018), which may contribute to the genomic variation observed for *Z. tritici* (Grandaubert *et al.* 2017; Hartmann *et al.* 2017). Under experimental conditions, the fungus has a narrow host range infecting wheat and shows abortive infections on closely related non-host grass species like *Triticum monococcum* (Jing *et al.* 2008) and *Brachypodium distachyon* (Kellner *et al.* 2014; O’Driscoll *et al.* 2015). However, the underlying determinants of host specialisation and host specificity of *Z. tritici* are largely unknown.

A previous study comparing the expression profiles of *Z. tritici* between early infection (4 days post infection) of the compatible host *T. aestivum* and the non-host *B. distachyon* revealed a set of 289 genes that were similarly expressed in the two hosts, but differentially expressed compared to growth in axenic culture (Kellner *et al.* 2014). These genes are likely crucial for *Z. tritici* during stomatal penetration that occurs in same way in both hosts. However, 40 genes showed differential expression between host and non-host infections (Kellner *et al.* 2014) and are possibly involved in the discrimination of compatible and non-compatible host-pathogen interactions. The signalling and regulatory networks responsible for these differential expression patterns are however unknown. One of the differentially expressed genes encodes the putative transcription factor Zt107320. Expression of *Zt107320* was significantly increased during infection of *T. aestivum* compared to the early infection of *B. distachyon* (Kellner *et al.* 2014) suggesting a host-dependent regulation of the gene.

*Zt107320* encodes a putative transcription factor belonging to the Zn(II)_2_Cys_6_ family. This gene family of transcription factors is exclusive to fungi (MacPherson *et al.* 2006; Pan and Coleman, 1990) and many members play an important role in the regulation of fungal physiology. For example, Zn(II)_2_Cys_6_ transcription factors in *Magnaporthe oryzae, Fusarium oxysporum, Leptosphaeria maculans, Parastagonospora nodorum* and *Pyrenophora tritici-repentis* are involved in the regulation of fungal growth and pathogenicity (Fox *et al.* 2008; Galhano *et al.* 2017; Imazaki *et al.* 2007; Lu *et al.* 2014; Rybak *et al.* 2017). Interestingly, the homolog of *Zt107320* in the rice blast pathogen *M. oryzae*, MoCOD1, was shown to affect conidiation and pathogenicity (Chung *et al.* 2013). *MoCOD1* was found to be upregulated during conidiation and appressorium formation at 72h post infection. Furthermore, the deletion mutant Δ*MoCOD1* showed defects in conidial germination and appressorium formation. *In planta*, the mutant Δ*MoCOD1* was attenuated in extending growth from the first-invaded cells and caused markedly reduced symptoms when compared to the wildtype (Chung *et al.* 2013).

Transcription factors, in general, regulate expression by integrating various signalling pathways and represent interesting targets for dissecting causes and mechanisms of pathogenicity and host specificity. In *M. oryzae*, a systemic approach was applied to characterise all 104 members of the Zn(II)_2_Cys_6_ family of transcription factors. Of these, 61 were shown to be involved in fungal development and pathogenicity (Lu *et al.* 2014). Similarly, in the head blight causing fungus *Fusarium graminearum* 26% of 657 tested transcription factors had an effect on the tested phenotypes mycelial growth, conidia production and toxin production (Son *et al.* 2011). In summary, a number of transcription factors, which play important roles in host infection and pathogenesis, have been identified for several important crop pathogens (Chen *et al.* 2017; Okmen *et al.* 2014; Xiong *et al.* 2015; Zhang *et al.* 2018; Zhuang *et al.* 2016). In *Z. tritici* however, only few regulatory genes have been characterised. Recently, two transcription factors ZtWor1 (Mirzadi Gohari *et al.* 2014) and ZtVf1 (Mohammadi *et al.* 2017) were shown to be important regulators of development and virulence of *Z. tritici* during compatible infections of wheat, highlighting how transcription factors can be used to identify and dissect aspects of pathogenicity. ZtWor1 has been functionally characterised and appears to be involved in the cAMP-dependent pathway, upregulated during the initiation of colonization and involved in regulating effector genes (Mirzadi Gohari *et al.* 2014). ZtVf1, a transcription factor belonging to the C_2_-H_2_ subfamily is required for virulence and its deletion leads to lower pycnidia density within lesions. Decreased virulence appears to be due to a reduced penetration frequency and impaired pycnidia differentiation (Mohammadi *et al.* 2017)

Based on the close homology of *Zt107320* to *MoCOD1* and its significantly different expression profile during host and non-host infections, we hypothesized that Zt107320 plays an important role during early wheat infection of *Z. tritici*. Our results confirm that Zt107320 affects virulence of *Z. tritici* during compatible infections and regulates the dimorphic switch as well as growth rate and cell wall properties of this important fungal plant pathogen.

## Results

### Phylogenetic analysis of Zt107320

In order to determine the distribution of homologs of Zt107320 we performed a similarity search on the protein level among putative fungal transcription factors. Among the 30 best matches 17 are found among either plant pathogens or plant associated fungal organisms, however this association does not appear to be monophyletic (fig 1 a). Among the thirty best hits, the *M. oryzae* homolog MoCod1 and the *Alternaria brassicola* homolog AbPf2 have been functionally analysed. Deletion of the *AbPf2* resulted in non-pathogenic strains (Cho *et al.* 2013). In the wild-type expression of *AbPf2* decreased after initial colonization of host tissues and the authors conclude that AbPf1 regulates pathogenesis (Cho *et al.* 2013). Among these highly conserved putative transcription factors two protein domains are shared: Zn(2)-C6 fungal-type DNA binding domain superfamily (IRP036864) and the transcription factor domain (IRP007219), indicating a functional role of the putative transcription factors (see fig. 1 b). Zt1073120 is therefore homologous to several predicted transcription factors in other fungal species, including some that have been associated with pathogenicity in plant pathogenic species, indicating a similar regulatory role of this protein in *Z. tritici*.

**Figure 1.**
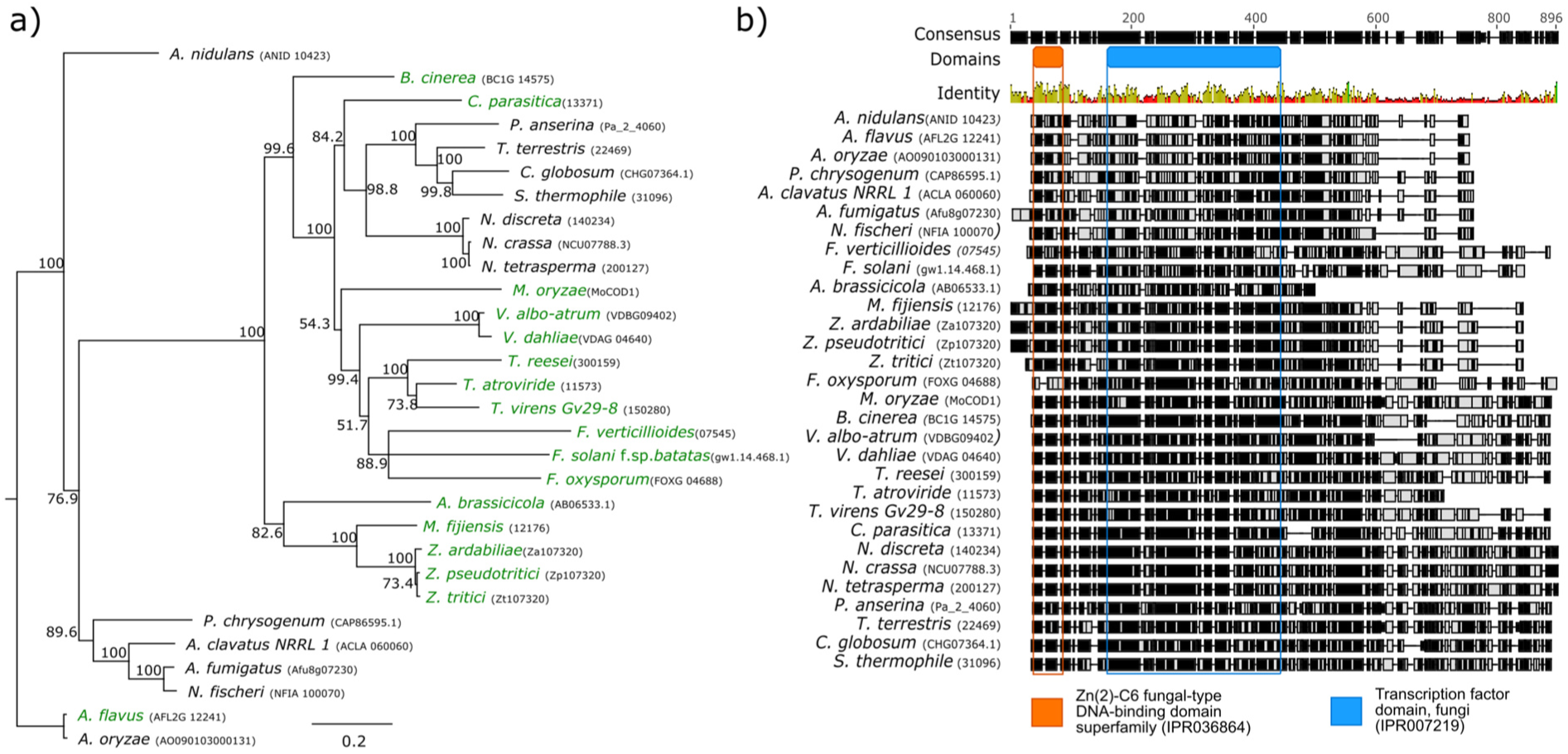
Homology of Zt107320 among fungal transcription factors. a) Phylogenetic tree based on 30 sequences showing the highest similarity with Zt107320 on the protein level, including the orthologs in the sister species *Z. pseudotritici* and *Z. ardabiliae*. The phylogenetic tree was constructed using MUSCLE alignments (Edgar, 2004) with Neighbour-Joining upon a consensus tree with 1000 bootstrapping iterations. Support of nodes by percentage of bootstrapping iterations is indicated. Species considered to be associated with plants are indicated in green. B) MUSCLE alignment of homologous sequences to Zt107320 indicating regions according to their identity with the consensus sequence. The localisations of two functional domains, identified using InterProScan, in the consensus sequence are indicated.

### Infections of *Z. tritici* are blocked in the substomatal cavities during incompatible interaction with *B. distachyon* coinciding with reduced expression of *Zt107320*

To study compatible and incompatible infections of *Z. tritici* in more detail, we inoculated leaves of 12-14 days old seedlings of *T. aestivum* (cultivar Obelisk) and *B. distachyon* (ecotype Bd21), and analysed the infection development by confocal microscopy. Fungal cells germinated upon contact with the leaf surface and developed infection hyphae. In contrast to previous observations (O’Driscoll *et al.* 2015), we found that *Z. tritici* infection hyphae entered into open *B. distachyon* stomata at four days post inoculation (dpi). However, further infection development of *Z. tritici* was blocked in the substomatal cavities of *B. distachyon* leaves (fig 2a), similar to phenotypes previously observed in incompatible interactions with einkorn wheat (Jing *et al.* 2008). Consequently, *Z. tritici* hyphae did not colonize the mesophyll tissue of *B. distachyon*, no necrotic lesions developed, and no asexual pycnidia formed. Interestingly, fungal growth was completely halted and no further growth was observed, even when leaves were examined five weeks post inoculation (fig 2a). In contrast, compatible infection of *Z. tritici* on the wheat cultivar Obelisk was characterised by previously described infection stages (Haueisen *et al.* 2019). Penetration of leaf stomata at 4 dpi was followed by the establishment of a hyphal network in the mesophyll tissue until 7-11 dpi and subsequently the switch to necrotrophy when biomass increased substantially resulting in the formation of pycnidia (fig 2a).

**Figure 2.**
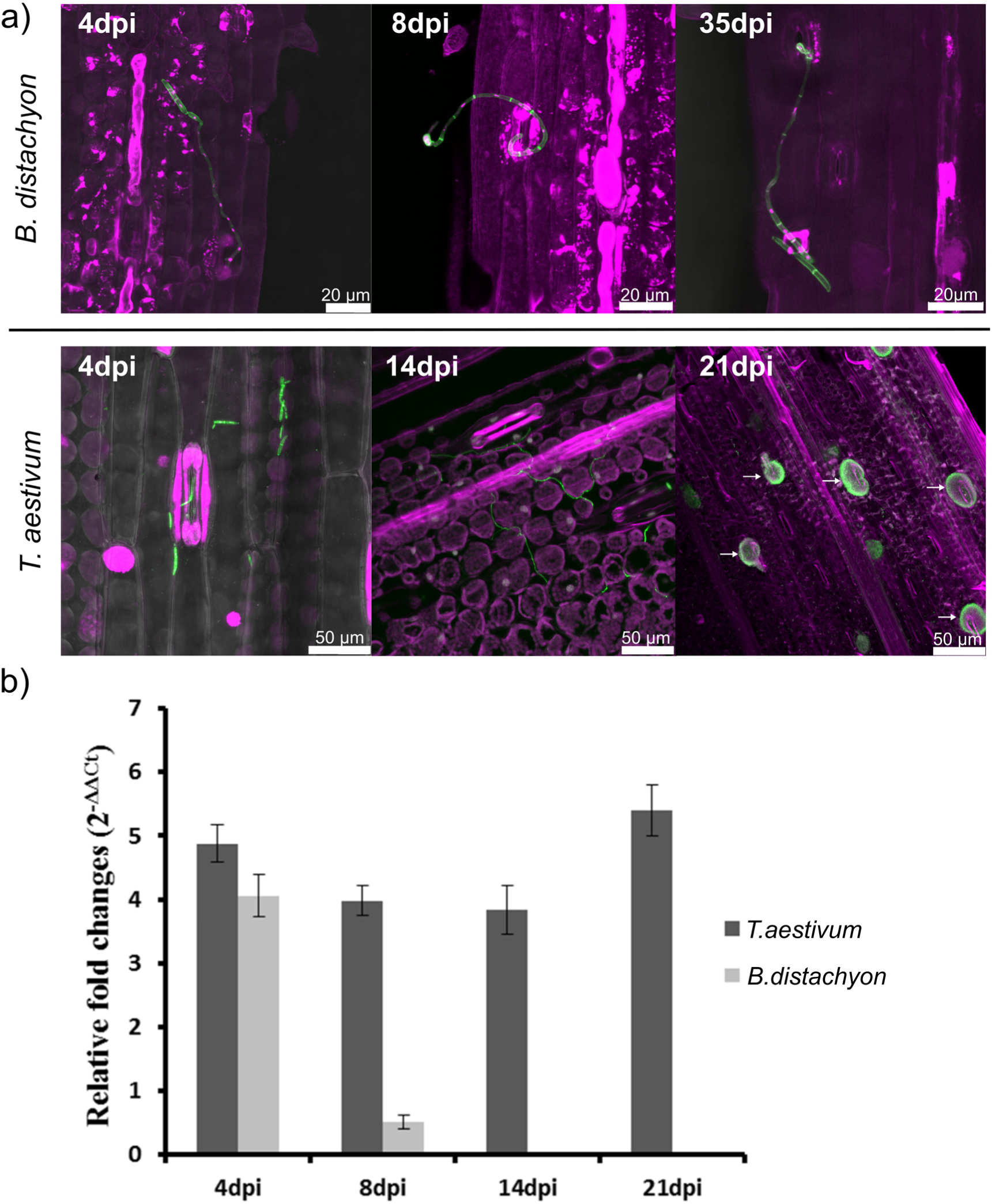
*Z. tritici* infection hyphae are blocked in the substomatal cavities of *B. distachyon* which coincides with reduced expression of *Zt107320*. a) Micrographs showing *Z. tritici* cells and emerging hyphae during penetration of *B. distachyon* stomata and infection of *T. aestivum*. On *T. aestivum, Z. tritici* germinated and penetrated the stomata at 4 dpi. Intercellular mesophyll colonization of *Z. tritici* occurred until 14 dpi resulting in the formation of pycnidia (indicated by arrows) at 21 dpi. In contrast, *Z. tritici* germinated on *B. distachyon* and penetrated stomata, but growth was halted below the stomata with no further leaf colonization (8 dpi - 35 dpi). Maximum projections of confocal image z-stacks. Nuclei and grass cells displayed in *purple* and fungal hyphae or septae respectively in *green*. b) Transcription of *Zt107320* in compatible (*T. aestivum)* and incompatible (*B. distachyon*) hosts. *In planta* expression of *Zt107320* is displayed relative to expression during axenic growth. Values are normalized to the expression of the gene encoding the housekeeping protein GAPDH. Error bars indicate the standard error of the mean (SEM) of three independent biological replicates per sample.

Previously, comparative transcriptome analyses during *Z. tritici* infection of the host *T. aestivum* and the non-host *B. distachyon* identified 40 differentially expressed genes at 4 dpi (Kellner *et al.* 2014). Building on this expression analysis, we focused on the putative transcription factor Zt107320 and validated the expression kinetics of *Zt107320* by RT-qPCR. We confirmed differential expression at 8 dpi (fig 2b). Expression in the non-host *B. distachyon* was greatly reduced at 8 dpi compared to expression in axenic cultures, whereas the expression of *Zt107320* during a compatible infection of the host *T. aestivum* was strongly upregulated during the course of the infection until 21 dpi.

### Zt107320 is located with the nucleus during yeast-like growth

The localization of Zt107320 within fungal cells was analysed using complementation strains that expressed a Zt107320_eGFP fusion protein regulated by the native promotor. During yeast-like growth on YMS media, the GFP-fusion protein appeared to be located in the nucleus or in its immediate vicinity (see fig 3). This localization is similar to the reported nuclear localization of the homolog AbPf1 (Cho *et al.* 2013) and supports the putative functional role of Zt107320 as a transcription factor. The observed localisation of the Zt107320_eGFP fusion protein also verifies the predicted nuclear localization of Zt107320 using the program WoLFPsort which implements an algorithm for prediction of subcellular locations of proteins based on sequence composition and content (Horton *et al.* 2007).

**Figure 3.**
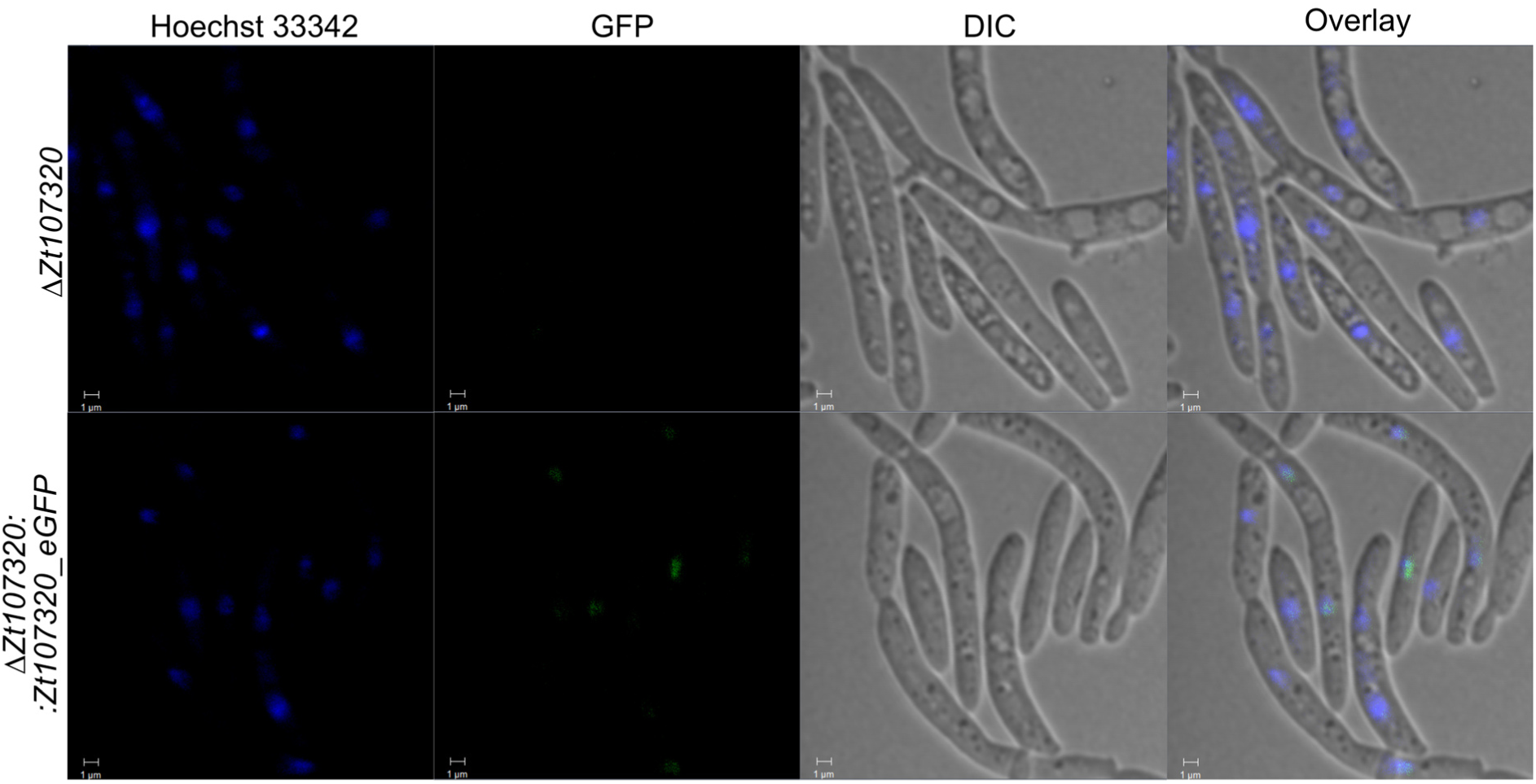
Localization of the Zt107320_eGFP fusion protein detected by fluorescence microscopy. Nuclei were counterstained using the DNA-specific dye Hoechst33342. eGFP fluorescence co-localized with the Hoechst33342 signal indicating nuclear localization of the Zt107320_eGFP fusion protein. eGFP fluorescence is restricted to the nuclei (scale bars = 1 µm).

### Zt107320 regulates growth and affects cell wall properties

Based on the coinciding halted growth and development and down-regulation of *Zt107320* in incompatible infections, we next asked whether Zt107320 is involved in the regulation of growth of *Z. tritici in vitro*. We determined the maximum growth rate r of *Z. tritici* in liquid cultures assuming a logistic growth curve model. For two independently generated *Zt107320* deletion mutants, we observed a significant reduction in the maximum growth rate compared to the wildtype. We thereby confirm the relevance of *Zt107320* as a putative regulator for growth, as two complementation strains showed no significant difference in the growth rate compared to the wildtype (fig 4a)

**Figure 4.**
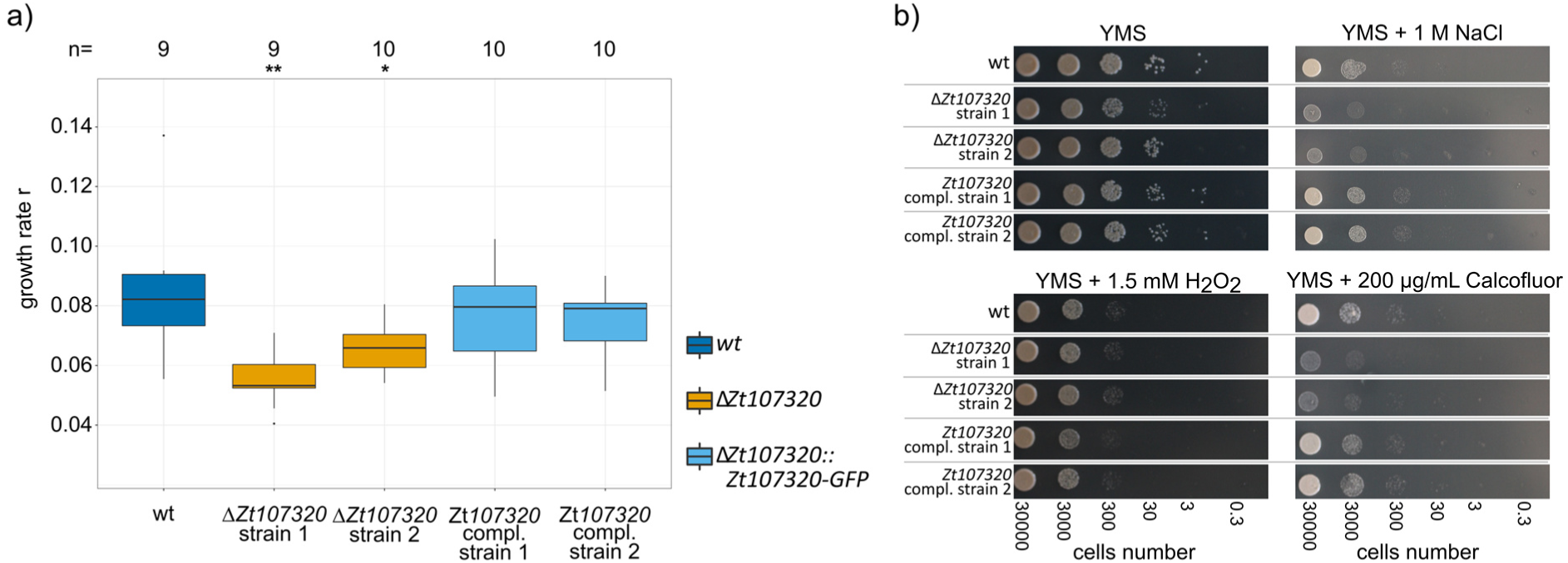
Zt107320 regulates growth and affects cell wall properties of *Z. tritici.* a) Maximum growth rate in liquid culture. Statistical significance inferred through an ANOVA and subsequent post hoc Tukey’s HSD comparing the deletion and complementation strains to the wildtype, is indicated as *= p <0.05; **= p<0.005; ***= p<0.0005. b) *In vitro* growth of the wildtype (wt), two deletion strains (Δ*Zt107320*), two complementation strains (Δ*Zt107320*::*Zt107320*_eGFP) on YMS media including the supplemented compounds to assess the role of osmotic stress (1 M NaCl), reactive oxygen species (1.5 mM H_2_O_2_) and cell wall stressors (200 µg/mL Calcofluor).

The effect of *Zt107320* deletion was not restricted to the growth rate but also included cell wall properties. Compared to the wildtype, we observed reduced growth *in vitro* when challenging the deletion mutants with high osmotic stresses (0.5 M and 1 M NaCl; 1 M, 1.5 M and 2 M Sorbitol) as well as with cell wall stress agents (300 μg/mL and 500 μg/mL Congo red; 200 μg/mL Calcofluor). Again, the wildtype phenotype was restored in both complementation strains (fig 4b and fig S1). Interestingly, temperature stress (28°C) did not affect the wildtype and the deletion mutants differently, as well as the exposure to H_2_O_2_ (1.5 mM and 2 mM). This indicates a specific effect of *Zt107320* on the cell wall properties but not on the ability of the fungus to counteract reactive oxygen species which are produced by the plant during activation of immune responses (Jones and Dangl, 2006).

The dimorphic switch from yeast-like to hyphal growth is considered to be central for pathogenicity during the early stages of infection (Kema, G. H. J. *et al.* 1996; Yemelin *et al.* 2017). We therefore next asked whether Zt107320 is involved in regulating this morphological switch of *Z. tritici.* To promote hyphal growth we used minimal medium and supplied different carbon sources in order to compare carbon utilisation and growth between the mutants and wildtype. Without carbon sources as well as in the presence of cellulose as exclusive carbon source, *Z. tritici* showed solely hyphal growth, indicating that carbon sources are required for yeast-like growth and that cellulose cannot be utilized by the fungus. The wildtype *Z. tritici* strain showed markedly increased yeast-like growth in the presence of the monosaccharides glucose, galactose, fructose and mannose as well as the sugar alcohols sorbitol and mannitol. Increased growth was also observed in the presence of the disaccharides sucrose and maltose (fig 5, fig S2). Interestingly, xylose as sole carbon source led to predominant hyphal growth in the wild type strain, which contrasts to the mainly yeast-like growth observed in the presence of all other tested carbohydrates. For xylose as sole carbon source, overall growth is markedly increased compared to the minimal medium lacking a carbon source indicating that xylose can be utilized by *Z. tritici* as a carbon source. The deletion of *Zt107320* affected fungal growth morphology on all tested carbon sources. Compared to the wildtype and the complementation strains, we observed increased hyphal growth for the Δ*Zt107320* strains on all carbon sources after 14 days of incubation, except for cellulose (fig 5, fig S2). In particular, supplementation of the monosaccharides fructose, glucose and xylose caused substantially pronounced hyphal growth when *Zt107320* was deleted. The wildtype phenotype was restored in both complementation strains confirming that *Zt107320* plays a role in the regulation of growth in *Z. tritici*.

**Figure 5.**
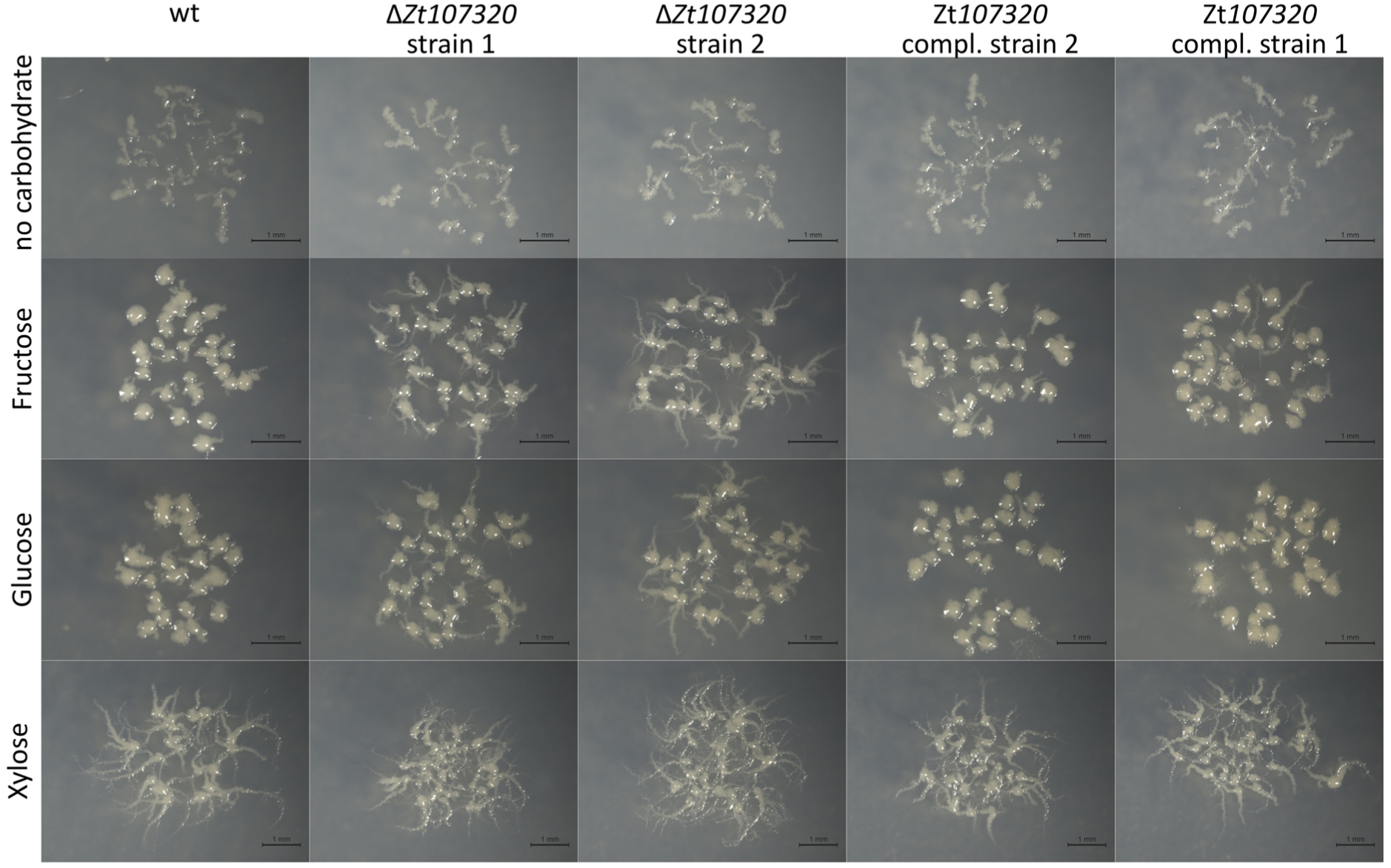
Zt107320 regulates the dimorphic switch of *Z. tritici*. Pictures showing morphologies of colonies originating from single cells after 14 days of growth at 18°C on minimal media containing the indicated carbohydrates as carbon sources. Fructose, glucose and xylose can be utilized by the fungus and led to increased growth. The two strains in which *Zt107320* was deleted independently showed an increase in hyphal growth compared to the wildtype, while in the two complementation strains the wildtype colony morphology was restored.

### Zt107320 affects the ability of *Z. tritici* to produce pycnidia

We next addressed whether the impact of *Zt107320* deletion on growth rate, the dimorphic switch and cell wall properties also influence the ability of *Z. tritici* to infect its host, *T. aestivum*. We inoculated a predefined area of the second leaf of the susceptible wheat cultivar Obelisk and measured the number of pycnidia 21 days post infection. We observed a pronounced reduction in the production of pycnidia for the *Zt107320* deletion strains. The density of pycnidia per cm^2^ of infected leaf area was reduced significantly for both independent deletions of *Zt107320* (ANOVA, p<1*10^-7^, p=0.002) (fig 6 a, b) compared to the wildtype. Complementing the *Zt107320* in-locus fully restored the wildtype phenotype *in planta*. As the density of pycnidia is considered to be a quantitative measure for virulence (Stewart *et al.* 2016), significantly decreased pycnidia production indicates reduced virulence of mutants. Thus, we conclude that the putative transcription factor Zt107320 by its effect on the growth rate, the dimorphic switch and the cell wall properties affects the fitness of *Z. tritici* during infection of the compatible wheat host.

**Figure 6.**
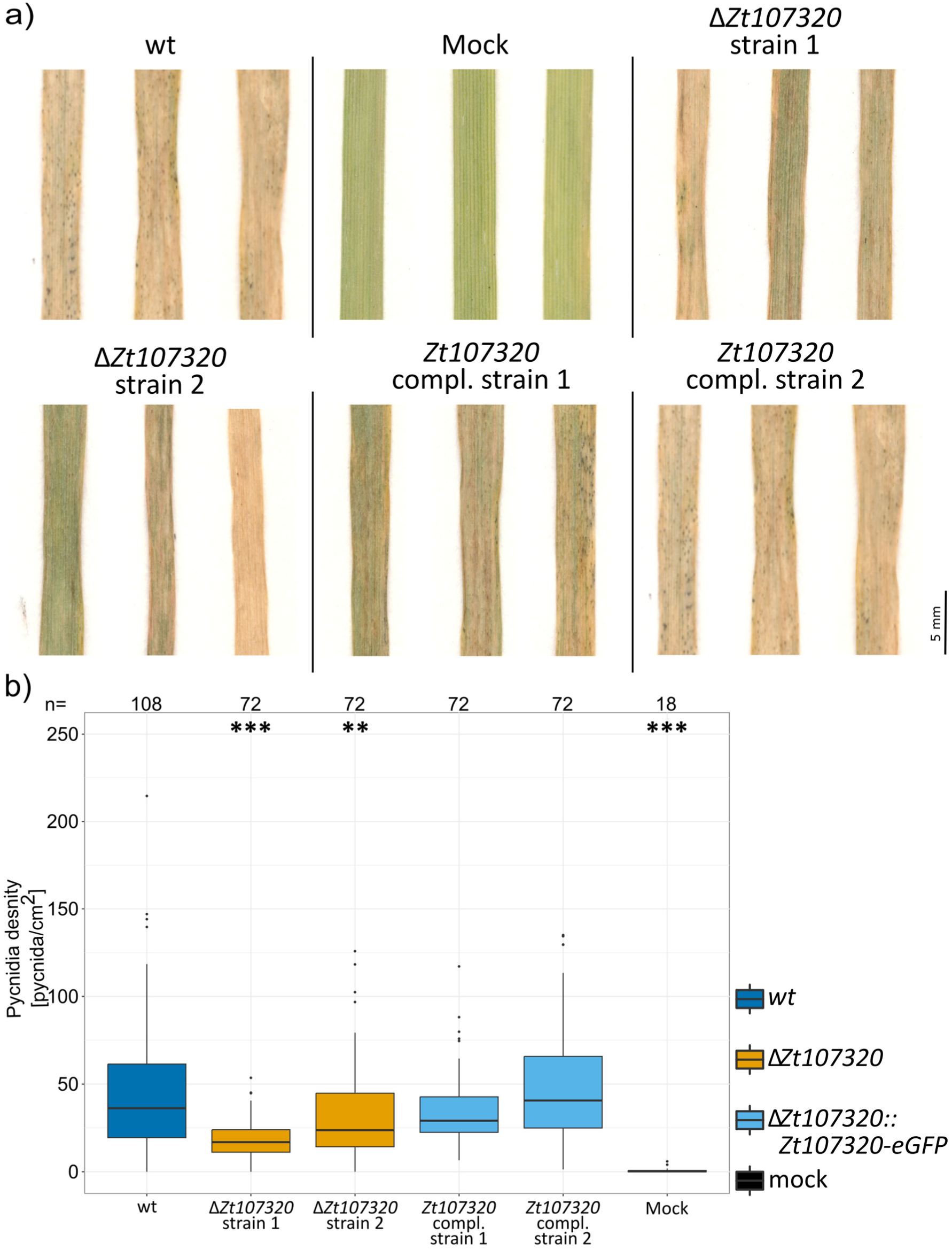
*Zt107320* deletion affects the pycnidia production during a compatible infection of wheat. a) Pictures of representative leaves infected with the *Z. tritici* wildtype (wt), two deletion strains (Δ*Zt107320*), two complementation strains (Δ*Zt107320*::*Zt107320*-*eGFP*) and mock-treated leaves. b) Boxplot depicting the pycnidia density (pycnidia per cm^2^ leaf) pooled for two independent experiments. Number of leaves (N) for each strain is indicated on top. Statistical significance, inferred through an ANOVA and subsequent post hoc Tukey’s HSD comparing the deletion and complementation strains to the wildtype, is indicated as *= p <0.05; **= p<0.005; ***= p<0.0005.

## Discussion

Here, we show that the putative transcription factor Zt107320, belonging to the fungal Zn(II)_2_Cys_6_ family, is involved in the infection program of *Z. tritici* during a compatible interaction with the host plant *T. aestivum*. Transcription of *Zt107320* is specifically induced during infection of wheat but not during the infection of the non-host *B. distachyon*. This suggests a functional role of this gene in the regulation of the infection program during a compatible host-pathogen interaction that allows the fungus to overcome host defences and propagate in the mesophyll tissue.

To date, only a small number of genes have been shown to be involved in virulence of this important wheat pathogen (Orton *et al.* 2011; Poppe *et al.* 2015). These genes encode the chitin binding LysM effector Mg3LysM (Lee *et al.* 2014; Marshall *et al.* 2011), two transcription factors ZtWor1 and ZtVf1 (Mirzadi Gohari *et al.* 2014; Mohammadi *et al.* 2017) and three rapidly evolving small proteins (Hartmann *et al.* 2017; Poppe *et al.* 2015; Zhong *et al.* 2017). Indeed, multiple effector candidate genes have been deleted but for most mutants no or small effects on pathogenicity were observed (Gohari *et al.* 2015; Rudd *et al.* 2015). Therefore, a high level of functional redundancy seems to be present in *Z. tritici*. The deletion of *Zt107320* results in a relatively small, nevertheless significant, reduction of the pycnidia density which is in contrast to the fully or partially avirulent phenotype demonstrated for the two previously deleted transcription factors ZtWor1 and ZtVf1, respectively (Mirzadi Gohari *et al.* 2014; Mohammadi *et al.* 2017). Similar to our observation in the *Zt107320* deletion mutant, deletion of *MoCOD1*, the rice blast homolog of *Zt107320*, led to a quantitative reduction in lesion size and number (Chung *et al.* 2013). This partial, but incomplete, quantitative reduction in virulence of *ΔZt107320* strains corresponds with the reduced, but still substantial growth rate of these deletion mutants *in vitro*. Therefore, other mechanisms, independent of Zt107320 are involved in the regulation of growth and development *in vitro* and *in planta*. Similarly, a partial, but incomplete reduction of invasive growth *in planta* as observed in Δ*MoCOD1* (Chung *et al.* 2013). The impact of Zt107320 on fungal growth and pathogenicity is therefore similar to the effect caused by other members of the Zn(II)_2_Cys_6_ transcription factor family: TPC1 regulates invasive, polarized growth and virulence in the rice blast fungus *M. oryzae* (Galhano *et al.* 2017), and the transcription factor FOW2 is known to control the ability of *F. oxysporum* to invade roots and colonize plant tissue (Imazaki *et al.* 2007).

Zt107320 appears to be involved in the morphological switch from yeast-like to hyphal growth as deletion of *Zt107320* led to an increase in hyphal growth. The switch to filamentous growth is considered to be essential for plant infection (Kema, G. H. J. *et al.* 1996; Yemelin *et al.* 2017). Therefore, the involvement of Zt107320 in regulating this dimorphism underscores its importance for pathogenicity. Based on RT-qPCR data we observe that *Zt107320* is differentially expressed between non-host and host infections and is further highly expressed during infections of compatible hosts. Other RNAseq-based transcriptome studies found that *Zt107320* is upregulated during later stages of wheat infection associated to necrotrophic host colonization, indicating a possible function for pycnidia formation (Haueisen *et al.* 2019; Rudd *et al.* 2015). Together, these findings support a role of Zt107320 in regulation of growth and pathogenicity of *Z. tritici*.

Interestingly, xylose the main product of hemicellulose degradation by fungal xylanases had a pronounced effect on the growth pattern of Z. *tritici*. Xylose supplementation not only increased overall growth compared to pure minimal medium but also led to an increase in the development of filaments spanning larger distances. In *Z. tritici*, the switch to necrotrophic growth after an initial phase of symptomless infection, overall disease severity and quantitative pycnidiospore production are associated with the activity of endo-β-1,4-xylanase (Siah *et al.* 2010; Somai-Jemmali *et al.* 2017). During the switch to necrotrophy *Z. tritici* rapidly develops large hyphal networks and uses plant-derived nutrients (Haueisen *et al.* 2019; Rudd *et al.* 2015). The observed effect of xylose on the growth morphology suggest that xylose – next to its role as a carbon source – may also direct growth and promote the spatial expansion of the intrafoliar hyphal network during the fungal lifestyle switch to necrotrophic growth. Indeed, genes encoding xylanases were shown to evolve under positive selection (Brunner *et al.* 2013), indicating an important functional role of this class of enzymes for adaptation to wheat infection. Although functional analyses of several xylanases in other plant pathogenic fungi, resulted in no direct phenotypic effect (Brunner *et al.* 2013; Douaiher *et al.* 2007) xylanases have been proposed as virulence factors (Douaiher *et al.* 2007). However, the results presented here indicate a possible role of xylose as a host infection-associated signal molecule for *Z. tritici* and should warrant further analysis.

In conclusion, we showed that Zt107320 affects the fitness of the wheat pathogen *Z. tritici*. *Zt107320* is differentially expressed in host and non-host environments, being upregulated during the early stages of infection on compatible hosts and down-regulated on non-hosts. This down-regulation corresponds to a considerably reduced growth and halted infections of *Z. tritici* after stomatal penetration of the non-host *B. distachyon*. In addition, we could confirm that Zt107320 has a nuclear localisation, consistent with its putative function as a transcription factor and that it further regulates the dimorphic switch between yeast-like and hyphal growth that is considered to be essential for pathogenicity. We therefore hypothesize that the putative transcription factor Zt107320 is part of to the regulatory network that controls host-associated growth and development, integrating signals that differ between compatible and non-compatible infections. Future studies should address the specific target genes of Zt107320 and their expression pattern during compatible and non-compatible interactions. Furthermore, the transcriptional regulation of *Zt107320* suggests that specific signals in the compatible host-pathogen interaction in wheat are responsible for the up-regulation of this particular transcription factor-encoding gene. Identification of these host-derived signals will provide fundamental insight into the molecular basis of host-pathogen interaction and host specificity in *Z. tritici*.

## Experimental Procedures

### Fungal and plant strains

The Dutch isolate IPO323 was kindly provided by Gert Kema (Wageningen, The Netherlands) and is available from the Westerwijk Institute (Utrecht, The Netherlands) with the accession number CBS 115943. The strain used in our experiments lacked accessory chromosome 18, presumably lost during culture maintenance *in vitro* (Kellner et al. 2014). Strains were maintained in either liquid Yeast Malt Sucrose (YMS) broth (4 g/L Yeast extract. 4 g/L Malt extract, 4 g/L sucrose) at 18°C on an orbital shaker or on solid YMS (+20 g/L agar) at 18°C. The *T. aestivum* cultivar Obelisk was obtained from Wiersum Plantbreeding BV (Winschoten, The Netherlands). *B. distachyon* inbred line Bd21 was kindly provided by Thierry Marcel (Bioger, INRA, France).

### Sequence analysis

Phylogenetic analysis of Zt107320 was conducted using the software Geneious Prime 2019.0.4 (https://www.geneious.com). Homologues sequences to protein sequence of Zt107320 were retrieved from the Fungal Transcription Factor Database (Park *et al.* 2008). Including Zt107320 and its homologs from the sister species *Z. pseudotritici* and *Z. ardabiliae* a total of the 30 best matches were retrieved. Alignments were constricted using MUSCLE (Edgar, 2004) and trees constructed using Neighbour-Joining algorithm building a consensus tree using 1000 bootstrapping replicates. Protein domains of the consensus sequence were identified using InterProScan (Quevillon *et al.* 2005). Prediction of the nuclear localisation of the Zt107320 was conducted using the WoLF PSORT predictor (Horton *et al.* 2007).

### Analysis of ***Z. tritici* during its compatible and non-compatible infections by confocal microscopy**

Morphology and development of *Z. tritici* inside and on the surface of leaves of B. *distachyon* inbred line Bd21 were analysed by confocal laser-scanning microscopy (CLSM) as described previously (Haueisen et al. 2019). Analyses of compatible *Z. tritici* infections on *Triticum aestivum* cultivar Obelisk were conducted by combining microtomy and CLSM as previously described (Rath *et al.* 2014). Distinct areas of the second leaf of 12-day-old (Bd21) and 14-day-old (wheat) seedlings were brush-inoculated with 1×10^7^ cells/ml in 0.1% Tween 20. Plants were incubated at 22°C [day]/ 20°C [night] and 100% humidity with a 16-h light period for 48 h. Subsequently, humidity was reduced to 70%. Microscopy was conducted using a Leica TCS SP5 and analysis of image z-stacks was done using Leica Application Suite Advanced Fluorescence (Leica Microsystems, Germany) and AMIRA® (FEITM Visualization Science Group, Germany).

### Analysis of Zt107320_eGFP expression using confocal microscopy

Cells were grown on solid YMS medium for 7 days before being scraped off the media surface and introduced into 10 mM phosphate buffer (pH 7.2) containing 1 μg/μl Hoechst 33342 (Sigma-Aldrich Chemie GmbH, Munich, Germany). Cells were incubated for 15-30 min in the dark and then transferred to a microscope slide and analysed by confocal laser scanning microscopy.

### RNA isolation and quantitative RT-PCR

We analysed gene expression patterns of *Zt107320* of *Z. tritici* employing a qRT-PCR experiment. Total RNA was extracted from fungal axenic cultures (grown for 72 h in YMS medium at 18°C and 200 rpm) and from snap-frozen leaf tissue infected with *Z. tritici* (4, 8, 14 and 21 dpi) using the TRIZOL reagent (Invitrogen, Karlsruhe, Germany), following the manufacturer’s instructions. Three biological replicates were included in the experimental set up. The cDNA samples were used in a qRT-PCR experiment employing the iQ SYBR Green Supermix Kit (Bio-Rad, Munich, Germany). PCR was conducted in a CFX96 RT-PCR Detection System (Bio-Rad, Munich, Germany) with the constitutively expressed control gene Glyceraldehyde-3-phosphate dehydrogenase (GAPDH). All primers are listed in table S1.

### Generation of Zt107320 deletion mutants and complementation by gene replacement

Fungal transformations and the creation of knock-out and complementation mutants of *Zt107320* were conducted as previously described (Poppe *et al.* 2015). In brief: Gene deletions were created by amplifying an approximately 1 kb region of the 5’ and 3’ flanking regions of *Zt107320* using PCR. The amplified flanking sequences were fused to a hygromycin resistance (hygR) cassette and an EcoRV cut vector backbone (pES61) using Gibson assembly (Gibson *et al.* 2009). Electro-competent cells of the *Agrobacterium tumefaciens* strains AGL1 were transformed using standard protocols. These transformed *A. tumefaciens* cells were used for the transformation of *Z. tritici* as previously described (Zwiers and De Waard, Maarten A. 2001). The same strategy was applied for the creation of the complementation strains by a C-terminal fusion of the *Zt107320* gene with an eGFP tag and a geneticin resistance cassette as a selection marker (fig. S3). After transformation and homologous recombination in the Δ*Zt107320* strain 1, the hygromycin resistance cassette was replaced by *Zt107320-eGFP* and the geneticin resistance cassette (NeoR) (fig. S3). Homologous recombination and integration was confirmed using PCR and Southern blot analysis by standard protocols. In short, genomic DNA was isolated using Phenol-Chloroform isolation (Sambrook and Russell, 2001). Restriction digestion was performed using PvuII, followed by gel-electrophoresis, blotting and detection using Dig-labelled probes binding to the upstream and downstream flank of *Zt107320* (fig. S2). In total four *ΔZt107320* strains, and two Δ*Zt107320*::*Zt107320*-eGFP strains were created.

### Isolation of fungal DNA and Southern blot analysis

DNA was isolated using phenol/chloroform applying a protocol previously described (Sambrook and Russell, 2001). Transformants were first screened using a PCR based approach detecting the resistance cassette and the endogenous locus. Candidate transformants were further confirmed by Southern blot analysis using a standard protocol (Southern, 1975). Probes were generated using the PCR DIG labelling Mix (Roche, Mannheim, Germany) according to the manufacturer’s instructions (Table S1).

### *In vitro* phenotyping

The *Z. tritici* strains were grown on YMS solid medium for 5-7 days at 18°C before the cells were scraped from the plate surface. For the determination of the growth rate, the cells were resuspended and counted. The cell density was adjusted to 50000 cells/mL and 175μl of the cell suspension was added to a well of a 96 well plate. Plates were inoculated at 18°C at 200 rpm with the OD600 being measured twice daily on a Multiskan Go plate reader (Thermo Scientific, Dreieich, Germany) (Table S2). Estimation of the growth rate was done by employing the logistic growth equation as implemented in the growthcurver package (version 0.2.1) in R (version R3.4.1) (R Core Team, 2015). For the determination of the *in vitro* phenotypes, the cell number was adjusted to 10^7^ cells/mL in ddH_2_O and serially diluted to 10^3^ cells/mL. 3 μL of each cell dilution was transferred onto YMS agar including the tested compounds and incubated for seven days at 18°C or 28°C. To test high osmotic stresses 0.5 M NaCl, 1 M NaCl, 1M Sorbitol, 1.5 M Sorbitol, 2 M Sorbitol (obtained from Carl Roth GmbH, Karlsruhe, Germany) were added to the YMS solid medium. To test cell wall stresses 300 μg/mL and 500 μg/mL Congo red and 200 μg/mL Calcofluor (obtained from Sigma-Aldrich Chemie GmbH, Munich, Germany) were added to the YMS solid medium. Finally, to determine the effect of reactive oxygen species on the mutant growth morphology 2 mM H_2_O_2_ (obtained from Carl Roth GmbH, Karlsruhe, Germany) was added to the YMS solid medium.

To test whether the Zt107320 affects the hyphal growth of *Z. tritici* and the ability of the fungus to use different carbon sources we used minimum media (MM) as described in (Barratt *et al.* 1965). Glucose, fructose, xylose, mannose, galactose, sorbitol, mannitol, sucrose and cellulose were added at a final concentration of 10 g/L and 20g/L agar were included. Strains were grown on YMS solid medium for 7 days and scraped into ddH2O, the cell number adjusted to 10^7^, 10^6^, 10^5^, 10^4^, 10^3^ cells/mL and 3 μl of each cell concentration added to the surface of the minimum medium plates. Plates were incubated 18°C in the dark and the growth monitored after 7 days and 14 days.

### *In planta* phenotyping

For the *in planta* phenotypic assays, we germinated seeds of the wheat cultivar Obelisk on wet sterile whatman paper for four days before potting using the soil Fruhstorfer Topferde (Hermann Meyer GmbH, Rellingen, Germany). Wheat seedlings were further grown for seven days before inoculation. *Z. tritici* strains were grown in YMS solid medium for five days at 18°C before the cells were scraped from the plate surface. The cell number was adjusted to 10^8^ cells/mL in H_2_O + 0.1% Tween 20, and the cell suspension was brushed onto approximately five cm on the abaxial and adaxial sides of the second leaf of each seedling. Inoculated plants were placed in sealed bags containing water for 48 h to facilitate infection through stomata. Plants were grown under constant conditions with a day night cycle of 16h light (~200μmol/m^2^*s) and 8h darkness in growth chambers at 20°C. Plants were grown for 21 days post inoculation at 90% relative humidity (RH). At 21 dpi the infected leaves were cut and taped to sheets of paper and pressed for five days at 4°C before being scanned at a resolution of 2400 dpi using a flatbed scanner (HP photosmart C4580, HP, Böblingen, Germany). Scanned images were analysed using an automated image analysis in Image J (Schneider *et al.* 2012) adapted from (Stewart *et al.* 2016). The read-out pycnidia/cm^2^ leaf surface was used for all subsequent analyses. See Table S3 for summary of *in planta* results.

### Statistical analysis

Statistical analyses were conducted in R (version R3.4.1) (R Core Team, 2015) using the suite R Studio (version 1.0.143) (RStudio Team, 2015). Data inspection showed a non-normal distribution for all data sets, including the measured pycnidia density (pycnidia/cm^2^). Therefore, we performed an omnibus analysis of variance using rank-transformation of the data (Conover and Iman, 1981) employing the model: pycnidia density ~ strain * experiment and r ~ strain, respectively. Post hoc tests were performed using Tukey’s HSD (Tukey, 1949).

## Acknowledgements

The study was funded by a personal grant to EHS from the State of Schleswig Holstein and a fellowship from the Max Planck Society, Germany. The funders had no role in study design, data collection and interpretation, or the decision to submit the work for publication. The authors have no conflict of interest.

## Supporting Information legends

**Figure S1. In vitro phenotype of *Zt107320* mutants.** I*n vitro* growth of the wildtype (wt), two independent deletion strains (Δ*Zt107320*), two independent complementation strains (Δ*Zt107320*::*Zt107320*_eGFP) on YMS media including the indicated compounds to assess the effect of osmotic stress (NaCl, Sorbitol), reactive oxygen species (H_2_O_2_), cell wall stressors (Calcofluor, Congo Red) and increased temperature (28°C) on growth and morphology of *Z. tritici.*

**Figure S2. Growth morphologies of Z. tritici wildtype and** Δ***Zt107320* and** Δ***Zt107320::Zt107320- eGFP* strains on minimum medium in the presence of different carbon sources.** Growth depicted after a) 7 days and b) 14 days. After 7 days, and more pronounced after 14 days of incubation, a higher amount of hyphal growth is observed for the Δ*Zt107320* mutants on the carbon sources Fructose, Galactose, Glucose, Maltose, Sorbitol, Sucrose and Xylose.

**Figure S3. Generation of *Zt107320* mutants in *Z. tritici***. a) Schematic illustration of the gene replacement strategy of *Zt107320*. Generation of the Δ*Zt107320* strains by homologous recombination between the upstream (UF) and downstream flanking regions (DF) of *Zt107320* in its genomic locus and a plasmid carrying the hygromycin resistance cassette (hygR) located between the UF and DF. Homologous recombination results in the integration of the hygromycin-resistance gene cassette (*hygR*) in the locus of *Zt107320*. Δ*Zt107320::Zt107300*-eGFP were generated by homologous recombination between UF and DF of the Δ*Zt107320* strains and a transformed plasmid containing a c-terminal fusion of *Zt107320* and *eGFP* and a Geneticin resistance cassette (*NeoR*) located between the UF and DF Yellow bars indicate the position of the probes used in the Southern blot analyses. b) Confirmation of *Zt107320* mutants by Southern blot analyses.

**Table S1: List of all primers used within this study**

**Table S2: Summary of *in vitro* growth data**

**Table S3: Summary of *in planta* phenotype**

