## Supplementary material for "The transcription factor Zt107320 affects the dimorphic switch, growth and virulence of the fungal wheat pathogen *Zymoseptoria tritici*": fig S1

- YMS
- 18°C

| Genotype | Strain |
|---|---|
| *ΔZt107320::Zt107320_eGFP* | Zt360#40 |
| *ΔZt107320::Zt107320_eGFP* | Zt360#5 |
| *ΔZt107320* | Zt356#38 |
| *ΔZt107320* | Zt356#20 |
| wt | Zt09 |
| *ΔZt107320::Zt107320_eGFP* | Zt360#40 |
| *ΔZt107320::Zt107320_eGFP* | Zt360#5 |
| *ΔZt107320* | Zt356#38 |
| *ΔZt107320* | Zt356#20 |
| wt | Zt09 |

30000 3000 300 30 3 0.3 cells

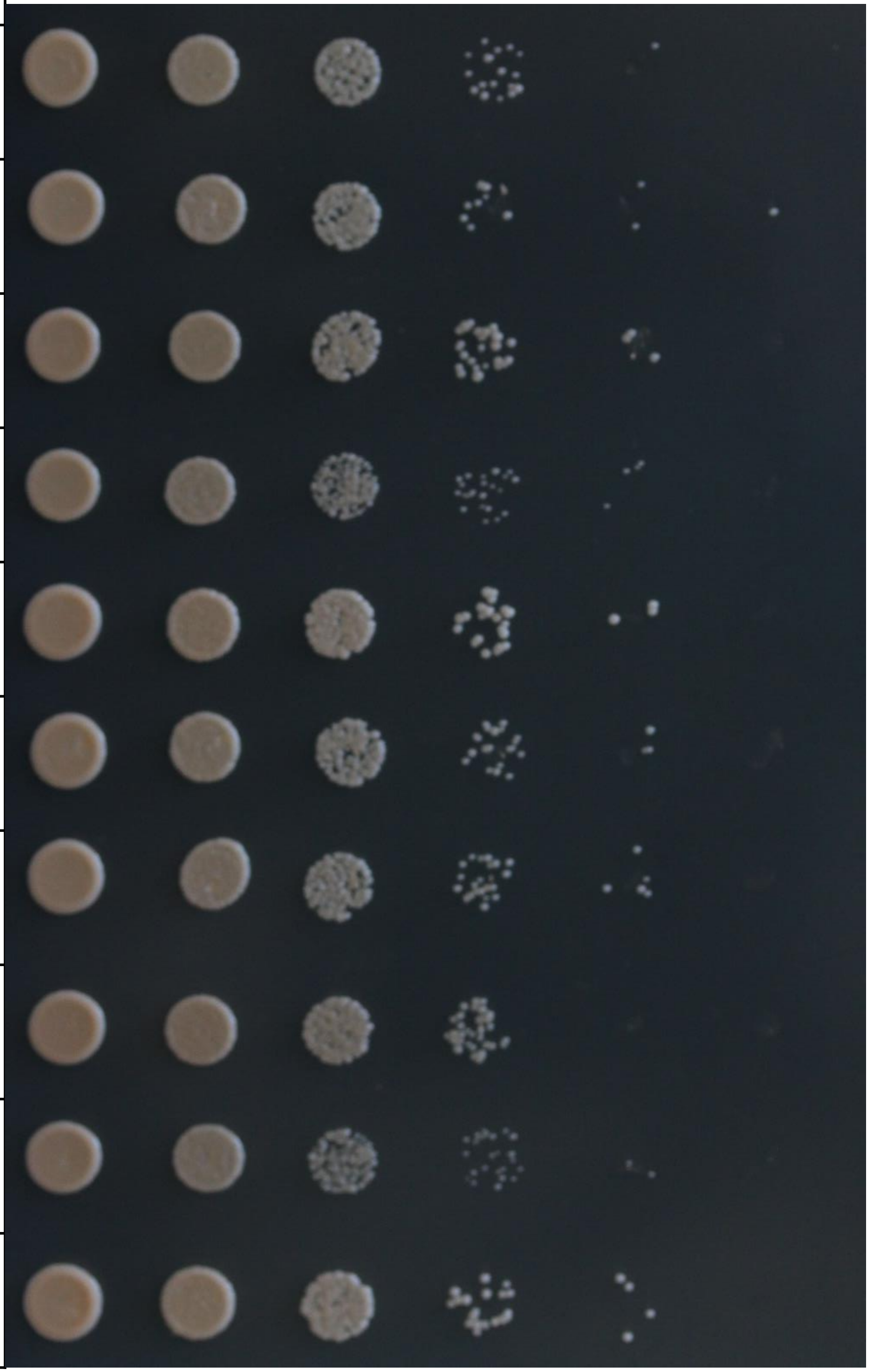

- YMS
- +1.5 mM $H_2O_2$
- 18°C

| Genotype | Strain |
|---|---|
| *ΔZt107320::Zt107320_eGFP* | Zt360#40 |
| *ΔZt107320::Zt107320_eGFP* | Zt360#5 |
| *ΔZt107320* | Zt356#38 |
| *ΔZt107320* | Zt356#20 |
| wt | Zt09 |
| *ΔZt107320::Zt107320_eGFP* | Zt360#40 |
| *ΔZt107320::Zt107320_eGFP* | Zt360#5 |
| *ΔZt107320* | Zt356#38 |
| *ΔZt107320* | Zt356#20 |
| wt | Zt09 |

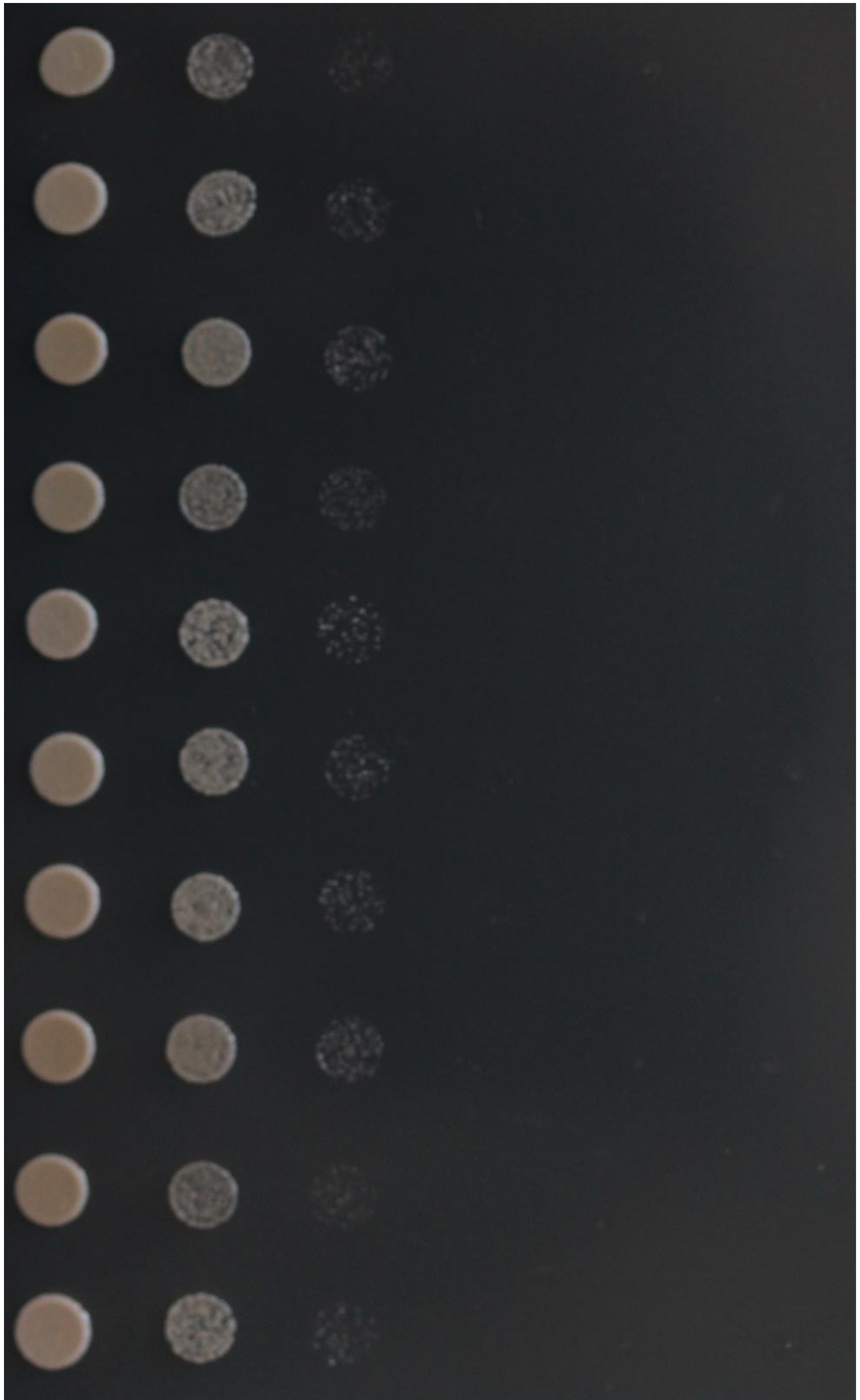

- YMS
- +2 mM $H_2O_2$
- 18°C

| Genotype | Strain |
|---|---|
| *ΔZt107320::Zt107320_eGFP* | Zt360#40 |
| *ΔZt107320::Zt107320_eGFP* | Zt360#5 |
| *ΔZt107320* | Zt356#38 |
| *ΔZt107320* | Zt356#20 |
| wt | Zt09 |
| *ΔZt107320::Zt107320_eGFP* | Zt360#40 |
| *ΔZt107320::Zt107320_eGFP* | Zt360#5 |
| *ΔZt107320* | Zt356#38 |
| *ΔZt107320* | Zt356#20 |
| wt | Zt09 |

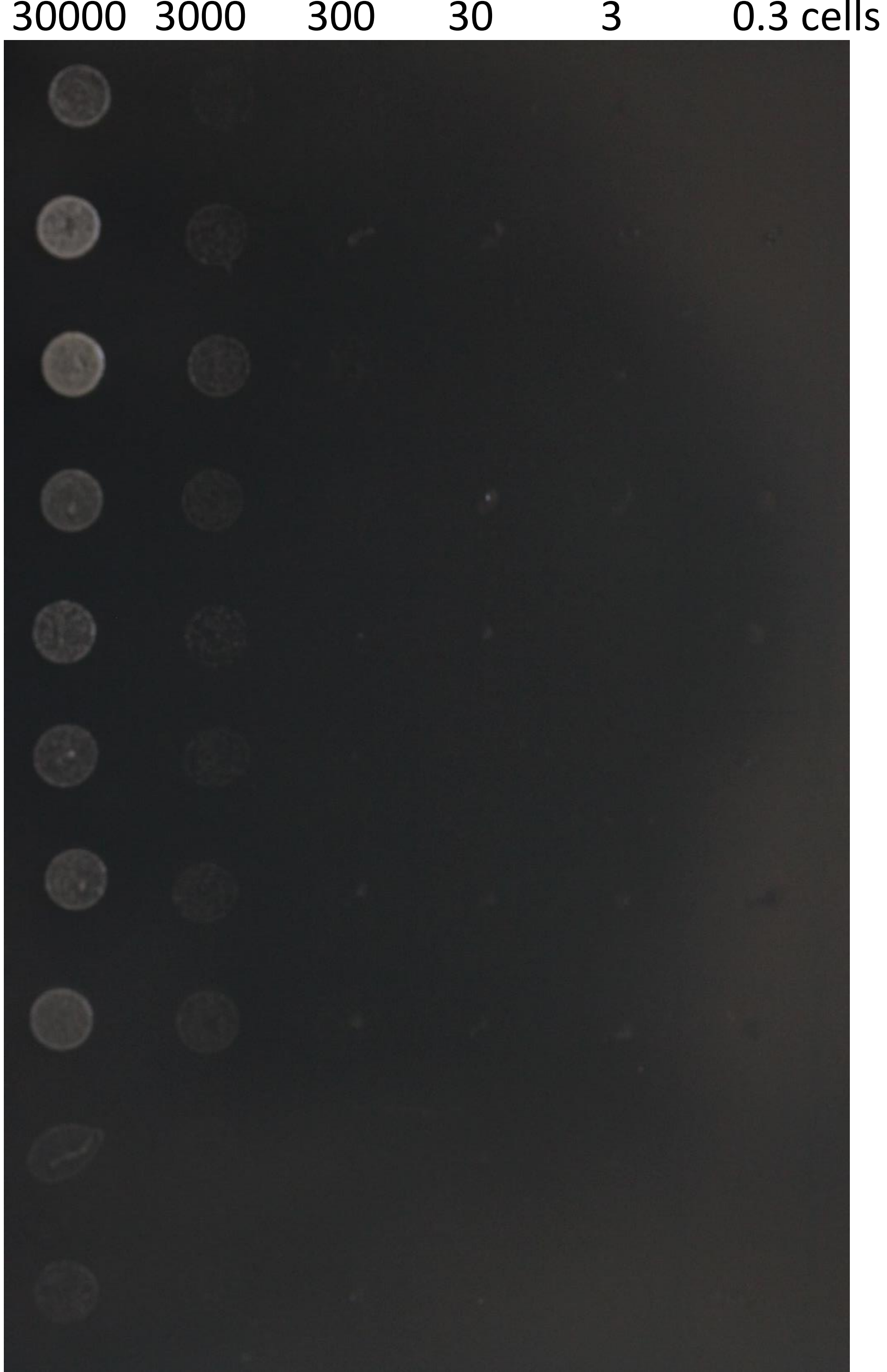

- YMS
- +500 µg/mL Congo red
- 18°C

| Genotype | Strain |
|---|---|
| *ΔZt107320::Zt107320_eGFP* | Zt360#40 |
| *ΔZt107320::Zt107320_eGFP* | Zt360#5 |
| *ΔZt107320* | Zt356#38 |
| *ΔZt107320* | Zt356#20 |
| wt | Zt09 |
| *ΔZt107320::Zt107320_eGFP* | Zt360#40 |
| *ΔZt107320::Zt107320_eGFP* | Zt360#5 |
| *ΔZt107320* | Zt356#38 |
| *ΔZt107320* | Zt356#20 |
| wt | Zt09 |

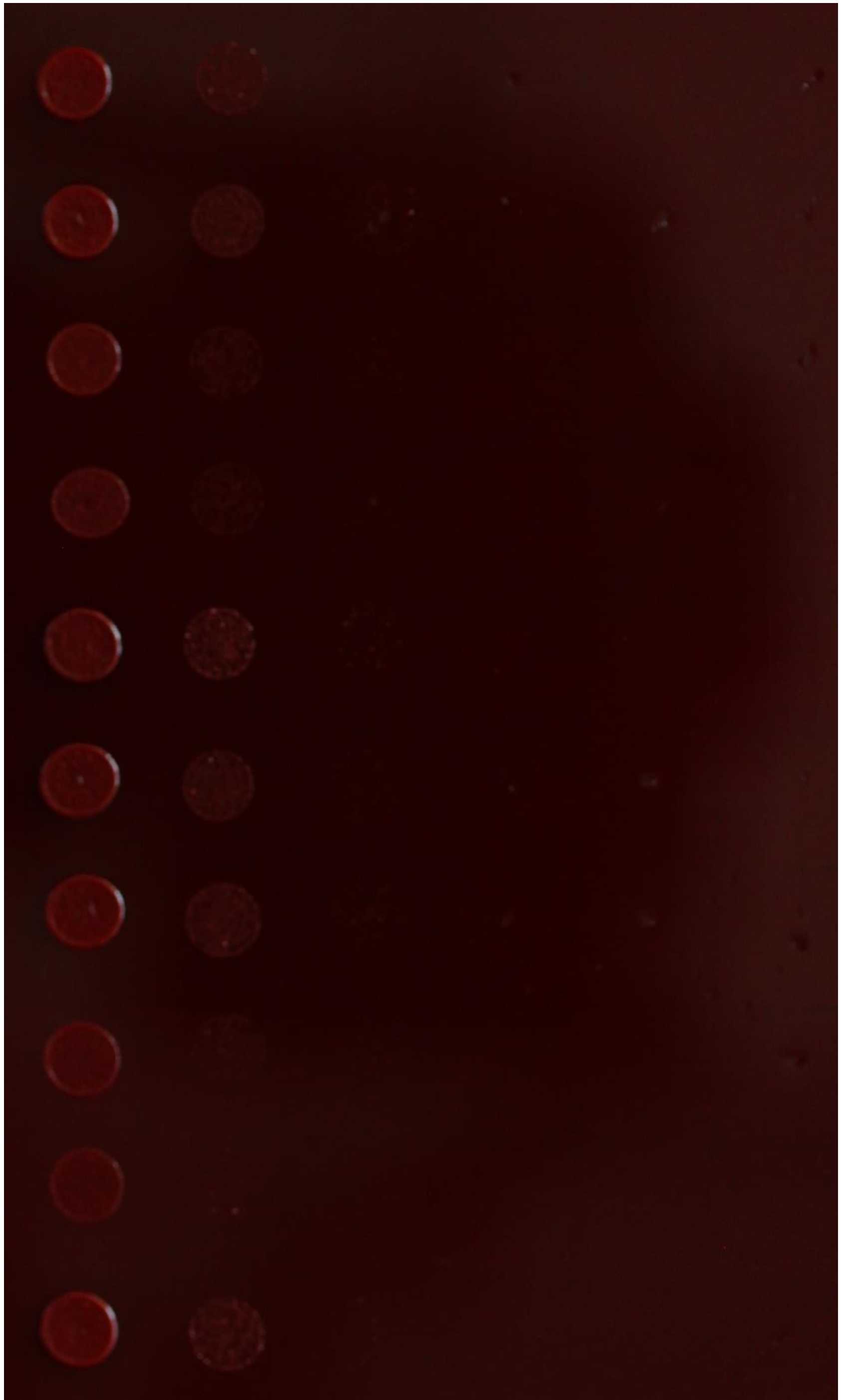

- YMS
- +300 µg/mL Congo Red
- 18°C

| Genotype | Strain |
|---|---|
| *ΔZt107320::Zt107320_eGFP* | Zt360#40 |
| *ΔZt107320::Zt107320_eGFP* | Zt360#5 |
| *ΔZt107320* | Zt356#38 |
| *ΔZt107320* | Zt356#20 |
| wt | Zt09 |
| *ΔZt107320::Zt107320_eGFP* | Zt360#40 |
| *ΔZt107320::Zt107320_eGFP* | Zt360#5 |
| *ΔZt107320* | Zt356#38 |
| *ΔZt107320* | Zt356#20 |
| wt | Zt09 |

30000 3000 300 30 3 0.3 cells

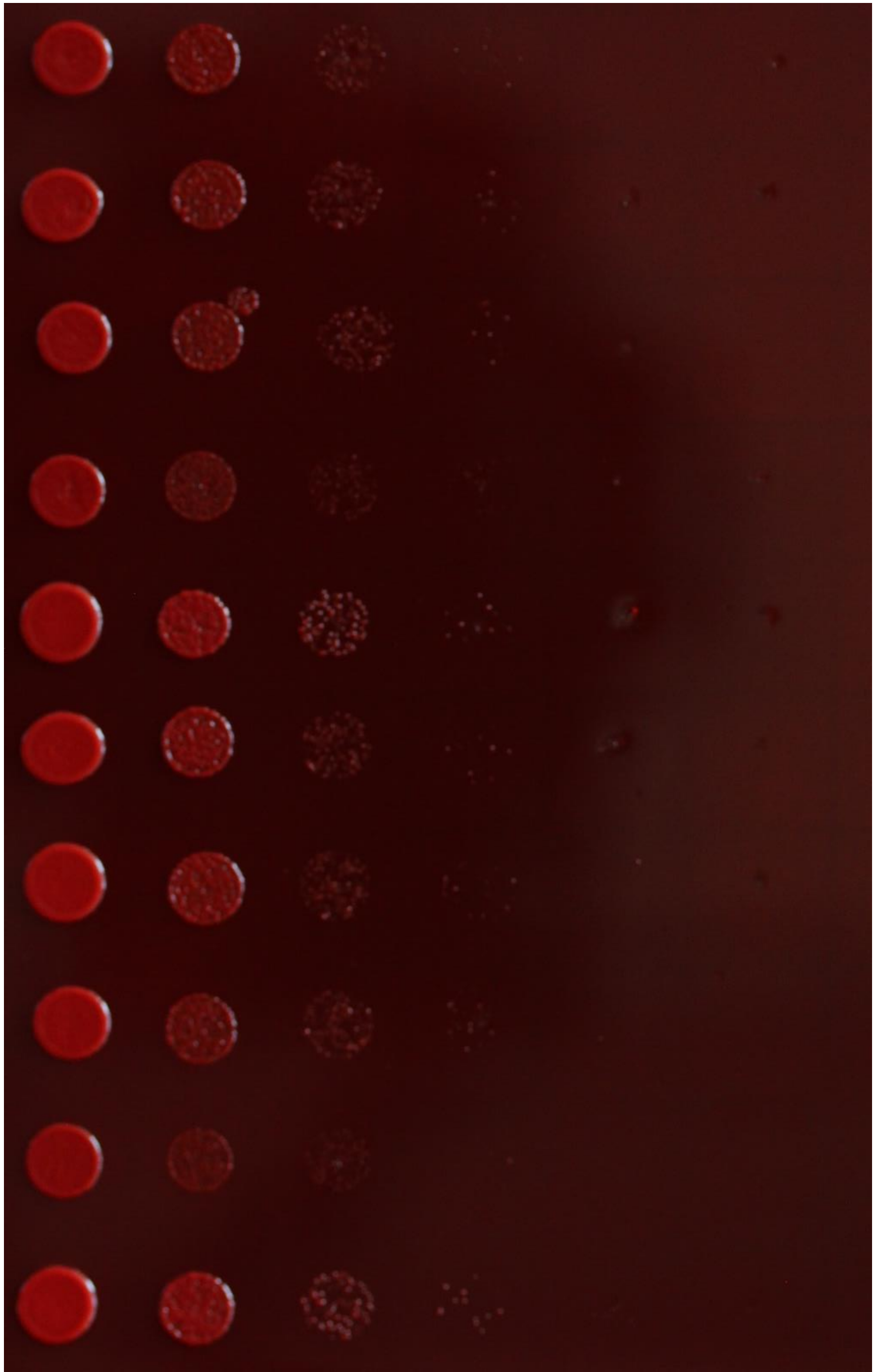

- YMS
- +200 µg/mL Calcofluor
- 18°C

| **Genotype** | **Strain** |
|---|---|
| *ΔZt107320::Zt107320_eGFP* | Zt360#40 |
| *ΔZt107320::Zt107320_eGFP* | Zt360#5 |
| *ΔZt107320* | Zt356#38 |
| *ΔZt107320* | Zt356#20 |
| wt | Zt09 |
| *ΔZt107320::Zt107320_eGFP* | Zt360#40 |
| *ΔZt107320::Zt107320_eGFP* | Zt360#5 |
| *ΔZt107320* | Zt356#38 |
| *ΔZt107320* | Zt356#20 |
| wt | Zt09 |

30000 3000 300 30 3 0.3 cells

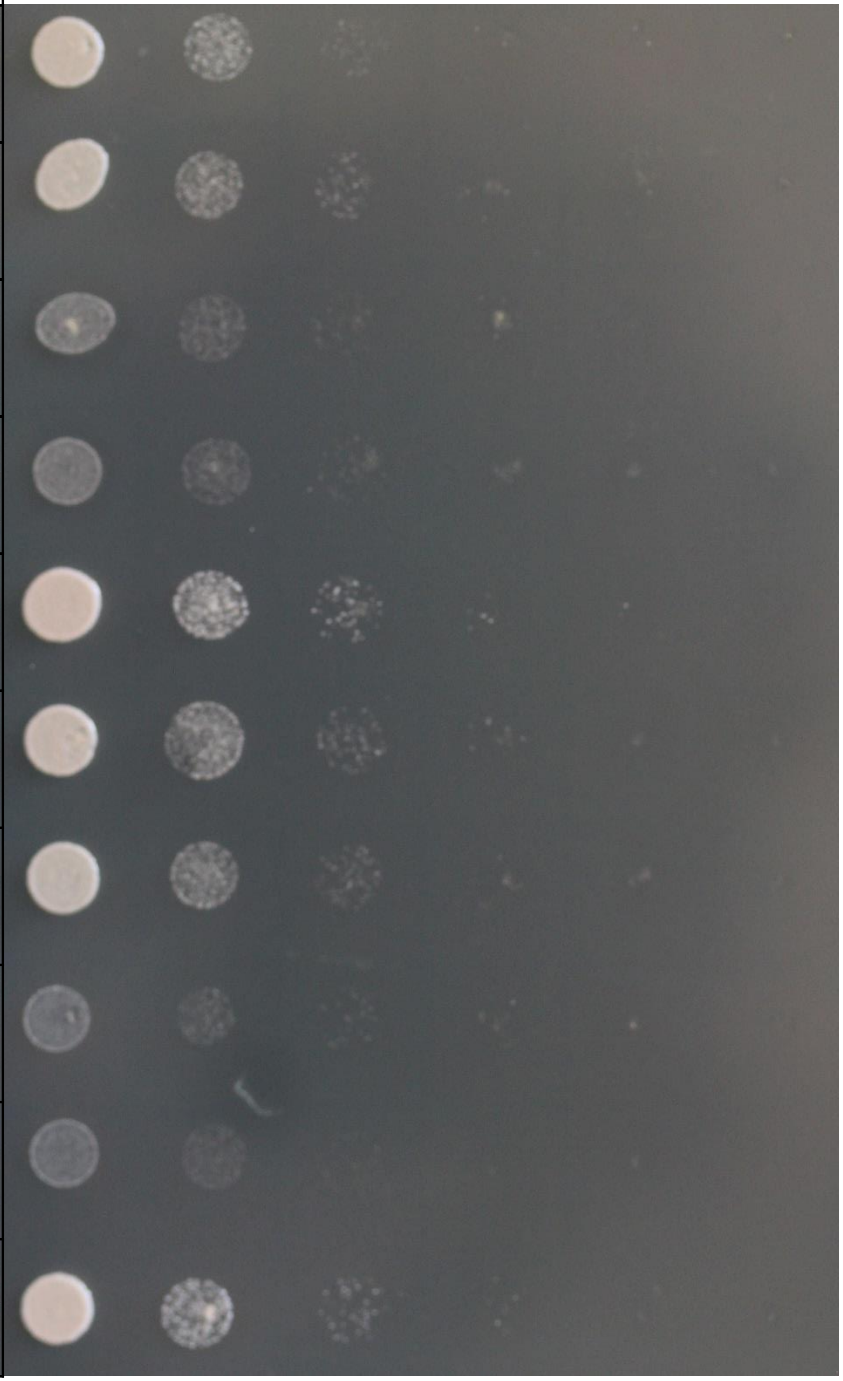

- YMS
- +0.5 M NaCl
- 18°C

| **Genotype** | **Strain** |
|---|---|
| *ΔZt107320::Zt107320_eGFP* | Zt360#40 |
| *ΔZt107320::Zt107320_eGFP* | Zt360#5 |
| *ΔZt107320* | Zt356#38 |
| *ΔZt107320* | Zt356#20 |
| wt | Zt09 |
| *ΔZt107320::Zt107320_eGFP* | Zt360#40 |
| *ΔZt107320::Zt107320_eGFP* | Zt360#5 |
| *ΔZt107320* | Zt356#38 |
| *ΔZt107320* | Zt356#20 |
| wt | Zt09 |

30000 3000 300 30 3 0.3 cells

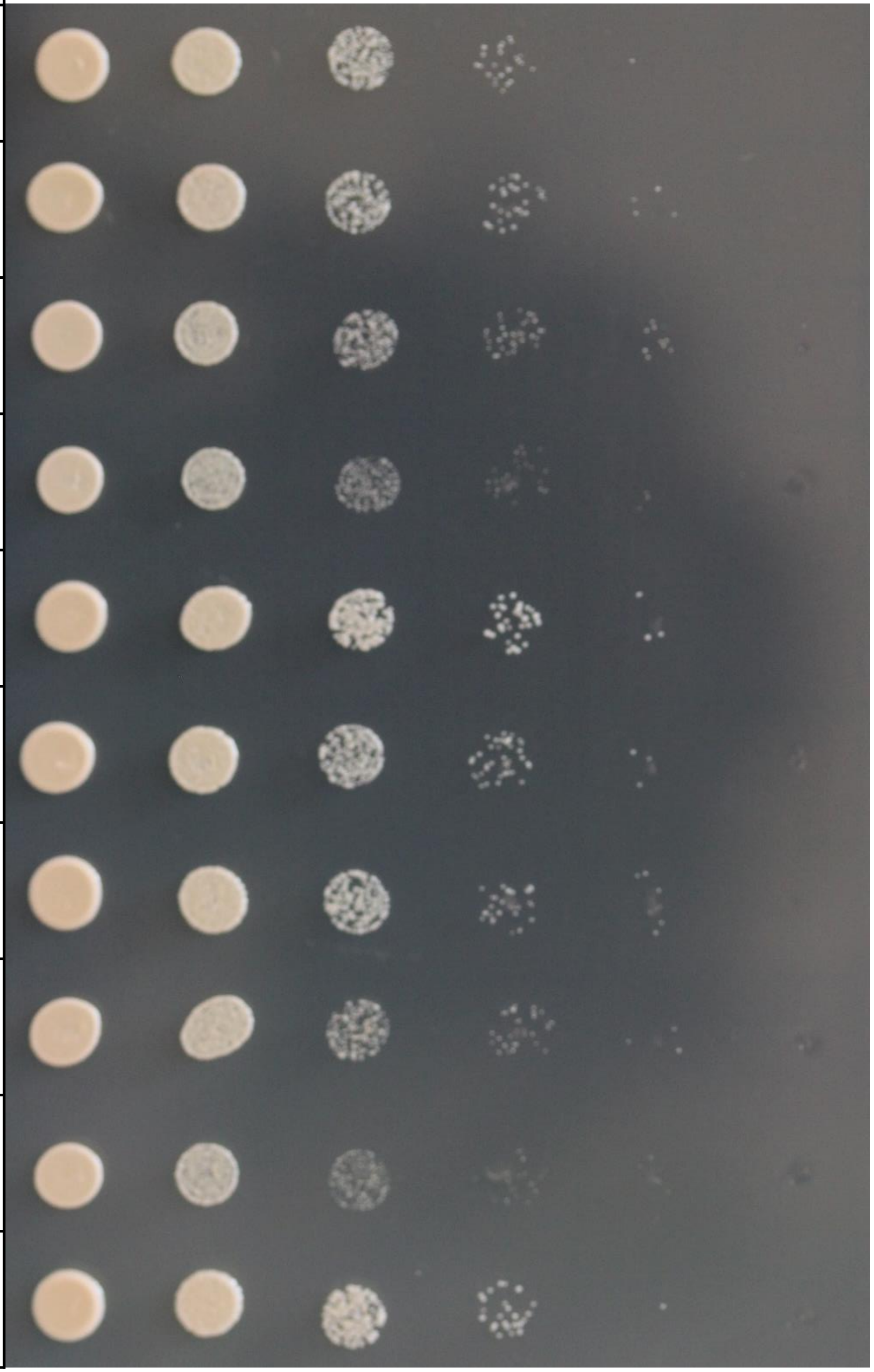

- YMS
- +1 M NaCl
- 18°C

| Genotype | Strain |
|---|---|
| *ΔZt107320::Zt107320_eGFP* | Zt360#40 |
| *ΔZt107320::Zt107320_eGFP* | Zt360#5 |
| *ΔZt107320* | Zt356#38 |
| *ΔZt107320* | Zt356#20 |
| wt | Zt09 |
| *ΔZt107320::Zt107320_eGFP* | Zt360#40 |
| *ΔZt107320::Zt107320_eGFP* | Zt360#5 |
| *ΔZt107320* | Zt356#38 |
| *ΔZt107320* | Zt356#20 |
| wt | Zt09 |

30000 3000 300 30 3 0.3 cells

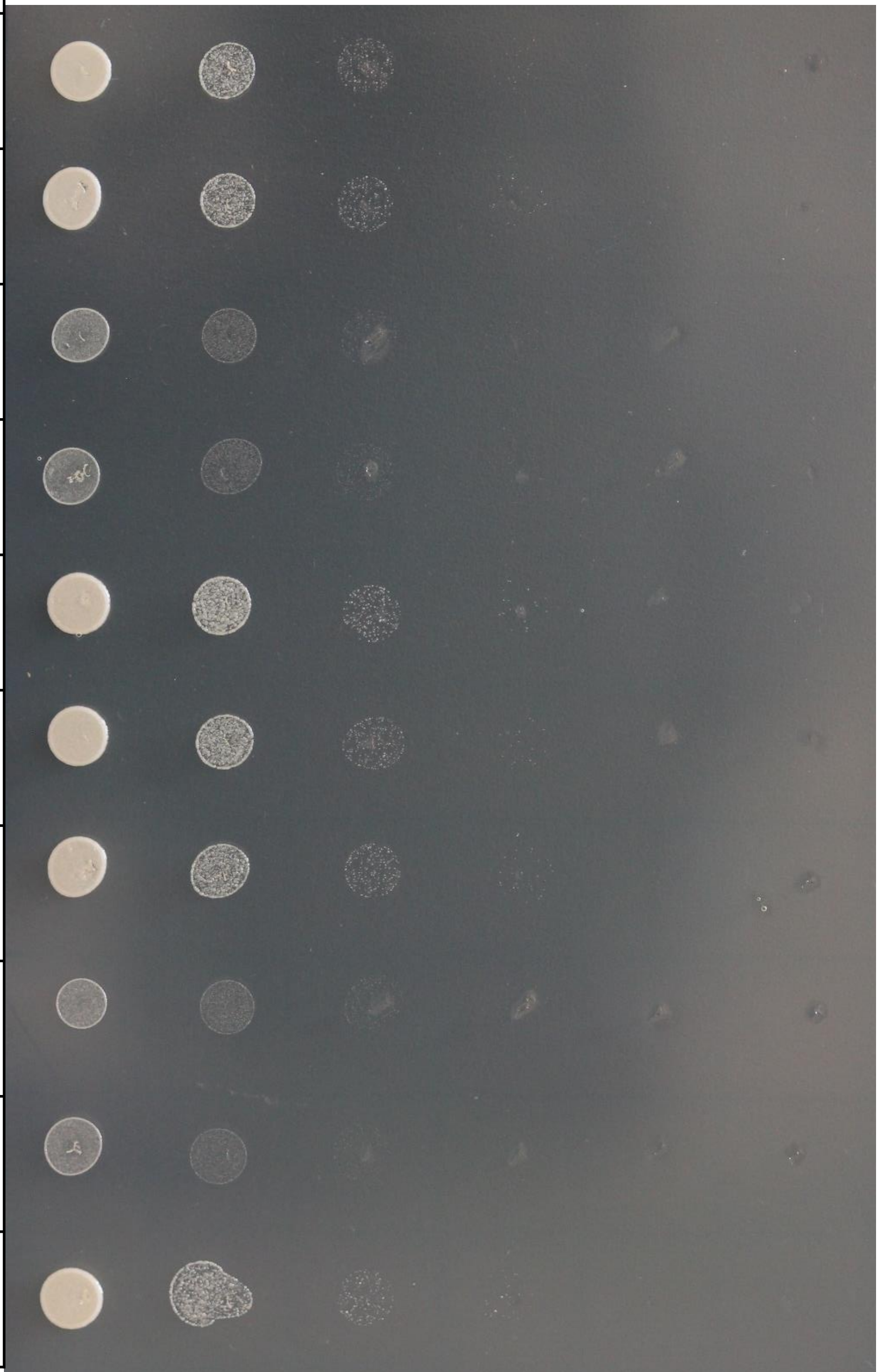

- YMS
- +1 M Sorbitol
- 18°C

| **Genotype** | **Strain** |
|---|---|
| *ΔZt107320::Zt107320_eGFP* | Zt360#40 |
| *ΔZt107320::Zt107320_eGFP* | Zt360#5 |
| *ΔZt107320* | Zt356#38 |
| *ΔZt107320* | Zt356#20 |
| wt | Zt09 |
| *ΔZt107320::Zt107320_eGFP* | Zt360#40 |
| *ΔZt107320::Zt107320_eGFP* | Zt360#5 |
| *ΔZt107320* | Zt356#38 |
| *ΔZt107320* | Zt356#20 |
| wt | Zt09 |

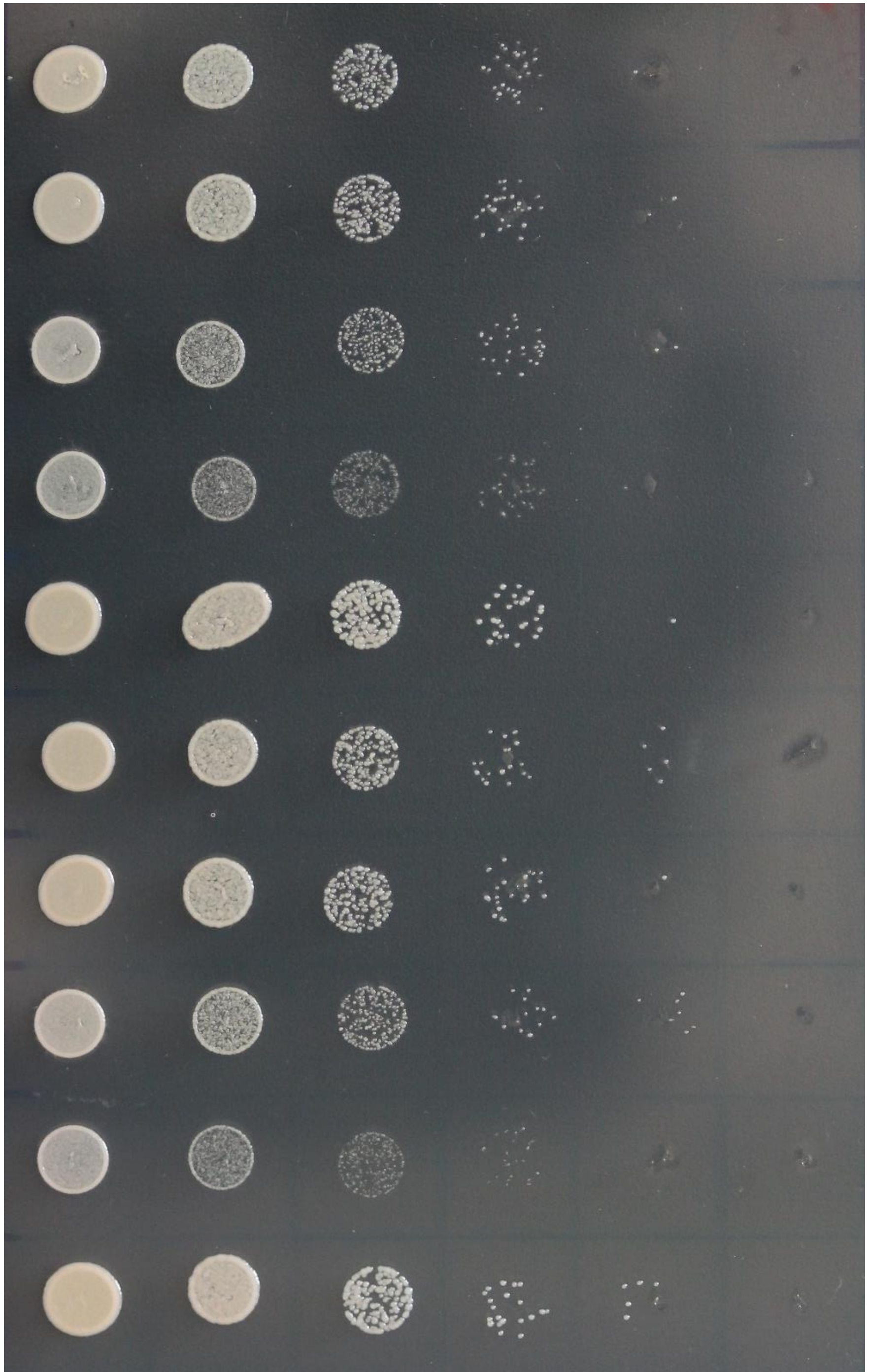

- YMS
- +1.5 M Sorbitol
- 18°C

| **Genotype** | **Strain** |
|---|---|
| *ΔZt107320::Zt107320_eGFP* | Zt360#40 |
| *ΔZt107320::Zt107320_eGFP* | Zt360#5 |
| *ΔZt107320* | Zt356#38 |
| *ΔZt107320* | Zt356#20 |
| wt | Zt09 |
| *ΔZt107320::Zt107320_eGFP* | Zt360#40 |
| *ΔZt107320::Zt107320_eGFP* | Zt360#5 |
| *ΔZt107320* | Zt356#38 |
| *ΔZt107320* | Zt356#20 |
| wt | Zt09 |

30000 3000 300 30 3 0.3 cells

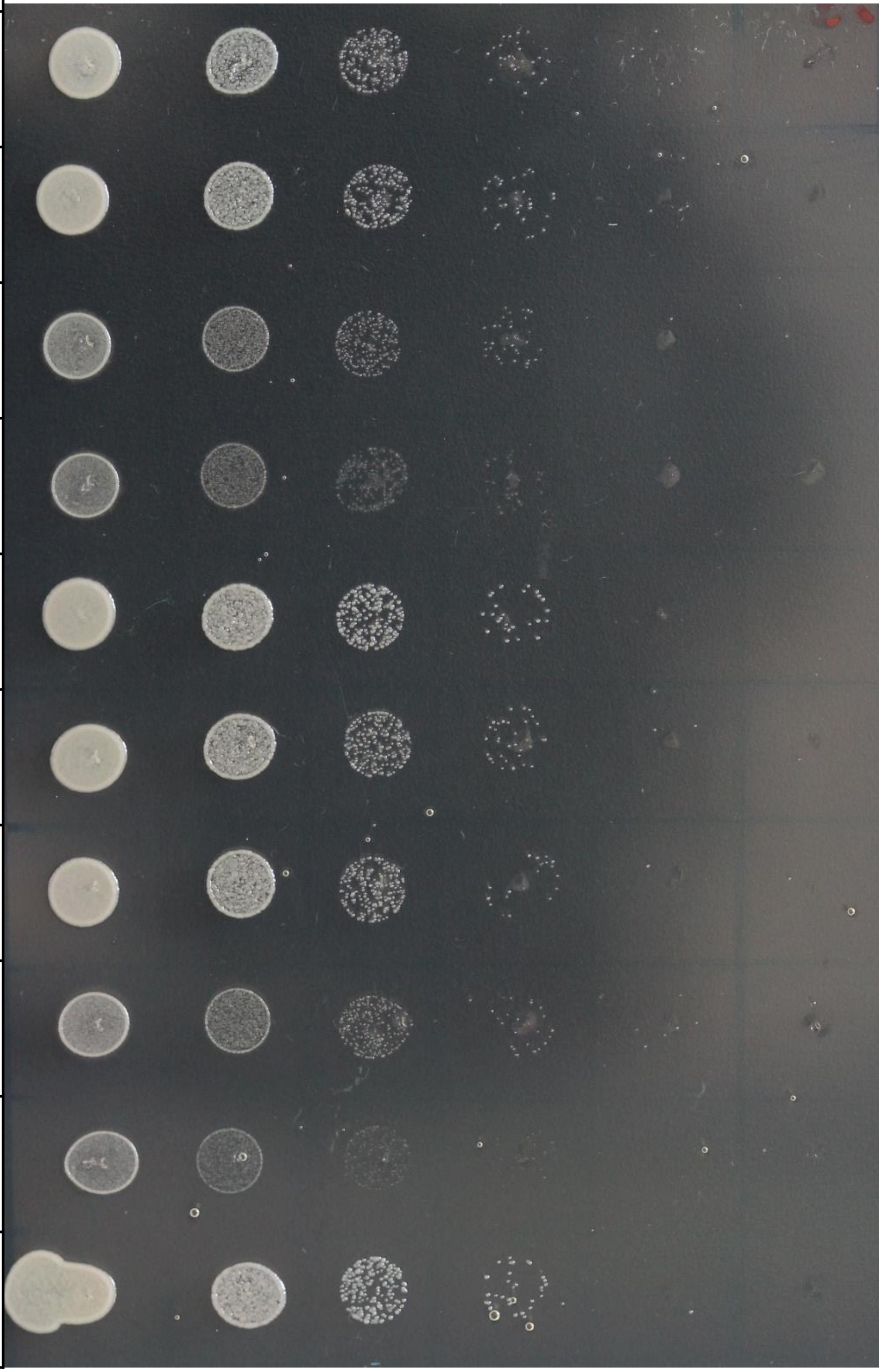

- YMS
- +2 M Sorbitol
- 18°C

| **Genotype** | **Strain** |
|---|---|
| *ΔZt107320::Zt107320_eGFP* | Zt360#40 |
| *ΔZt107320::Zt107320_eGFP* | Zt360#5 |
| *ΔZt107320* | Zt356#38 |
| *ΔZt107320* | Zt356#20 |
| wt | Zt09 |
| *ΔZt107320::Zt107320_eGFP* | Zt360#40 |
| *ΔZt107320::Zt107320_eGFP* | Zt360#5 |
| *ΔZt107320* | Zt356#38 |
| *ΔZt107320* | Zt356#20 |
| wt | Zt09 |

30000 3000 300 30 3 0.3 cells

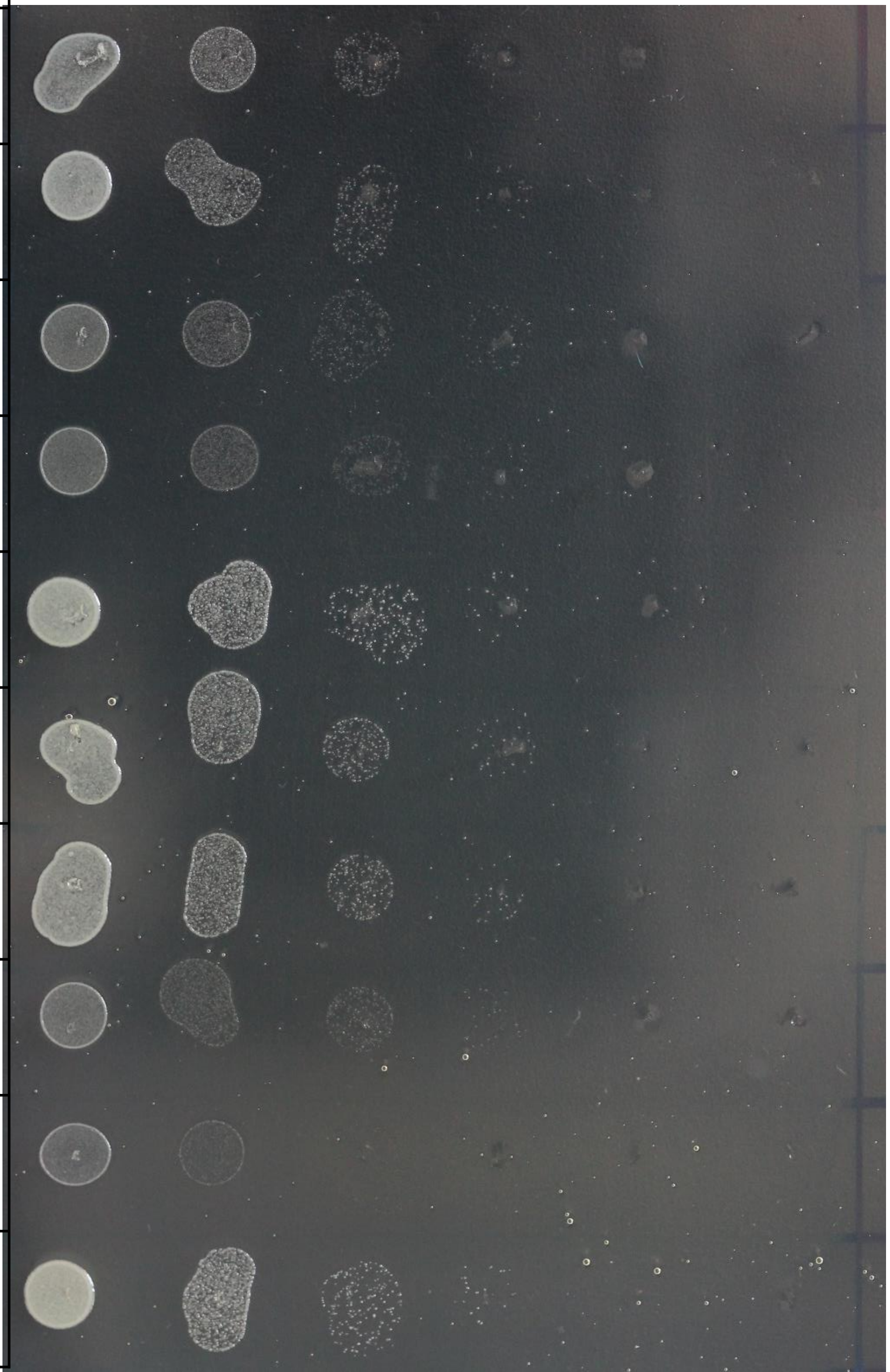

- YMS
- 28°C

| Genotype | Strain |
|---|---|
| *ΔZt107320::Zt107320_eGFP* | Zt360#40 |
| *ΔZt107320::Zt107320_eGFP* | Zt360#5 |
| *ΔZt107320* | Zt356#38 |
| *ΔZt107320* | Zt356#20 |
| wt | Zt09 |
| *ΔZt107320::Zt107320_eGFP* | Zt360#40 |
| *ΔZt107320::Zt107320_eGFP* | Zt360#5 |
| *ΔZt107320* | Zt356#38 |
| *ΔZt107320* | Zt356#20 |
| wt | Zt09 |

30000 3000 300 30 3 0.3 cells

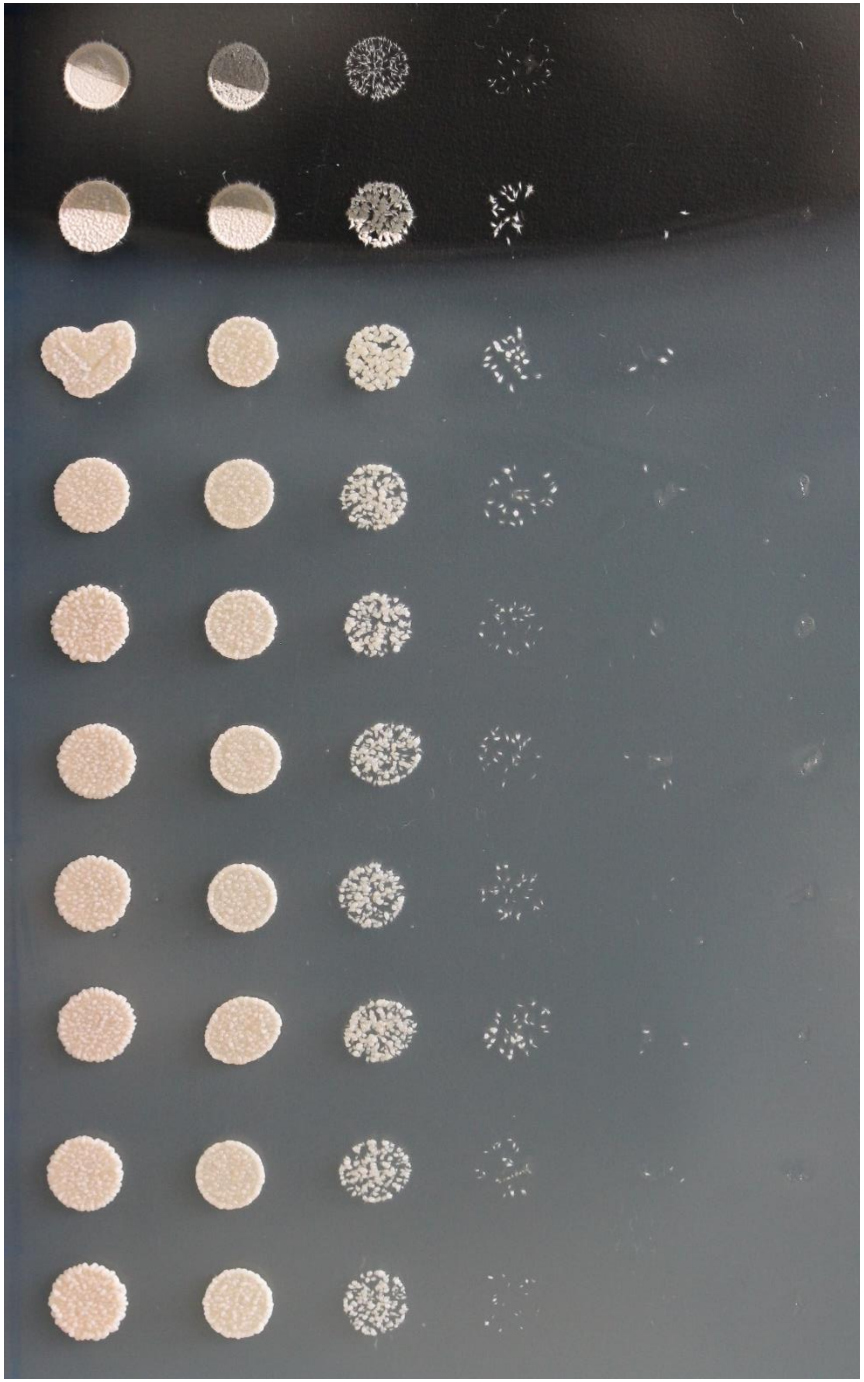

**Figure S1. In vitro phenotype of *Zt107320* mutants**

*In vitro* growth of the wildtype (wt), two independent deletion strains (Δ*Zt107320*), two independent complementation strains (Δ*Zt107320*::*Zt107320*_eGFP) on YMS media including the indicated compounds to assess the effect of osmotic stress (NaCl, Sorbitol) , reactive oxygen species ($H_2O_2$), cell wall stressors (Calcofluor, Congo Red) and increased temperature (28°C) on growth and morphology of *Z. tritici.*
