## Supplementary figures and images for "The transcription factor Zt107320 affects the dimorphic switch, growth and virulence of the fungal wheat pathogen *Zymoseptoria tritici*"

### fig S2

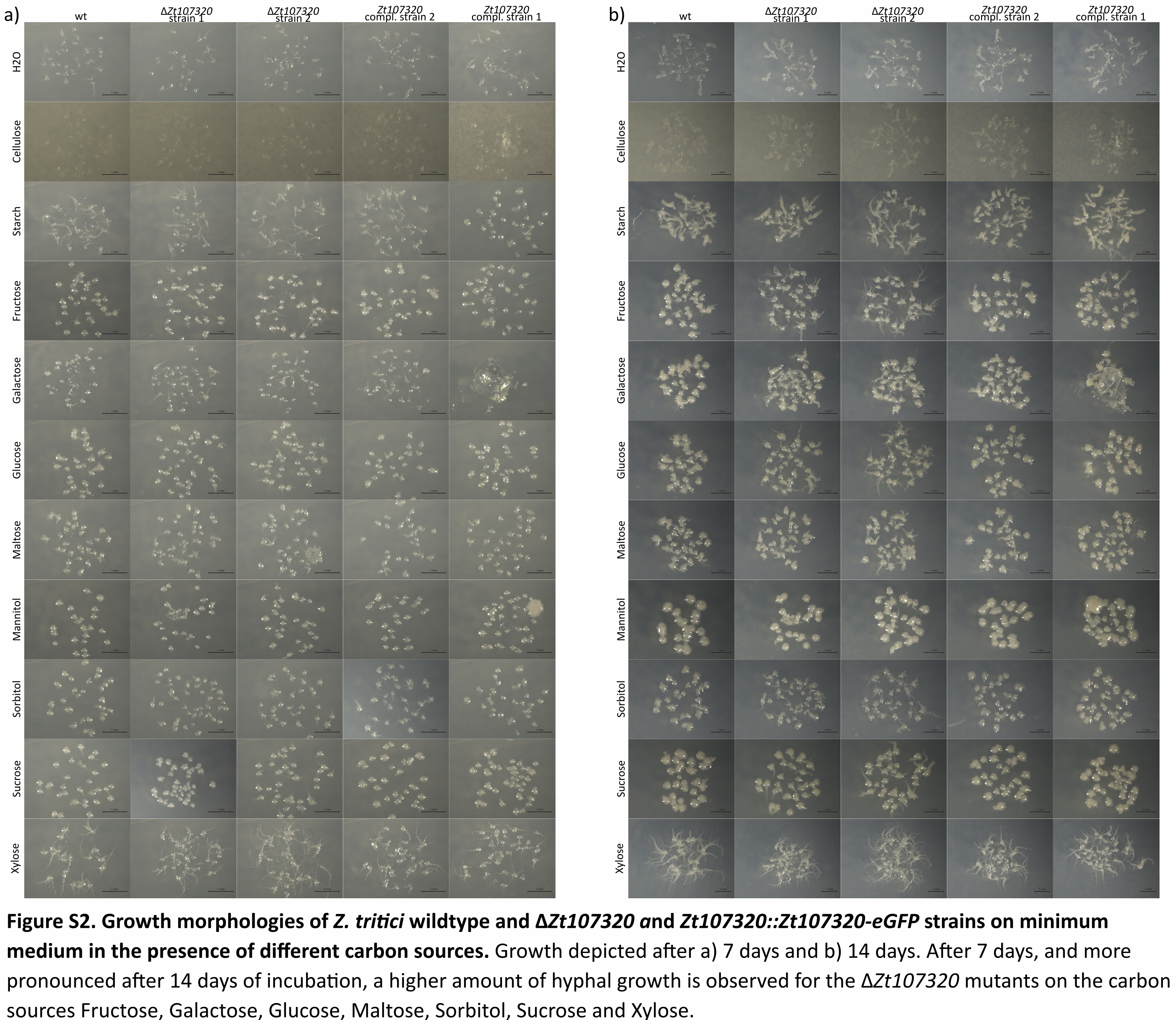
