## Supplementary material for "The transcription factor Zt107320 affects the dimorphic switch, growth and virulence of the fungal wheat pathogen *Zymoseptoria tritici*": fig S3

a)

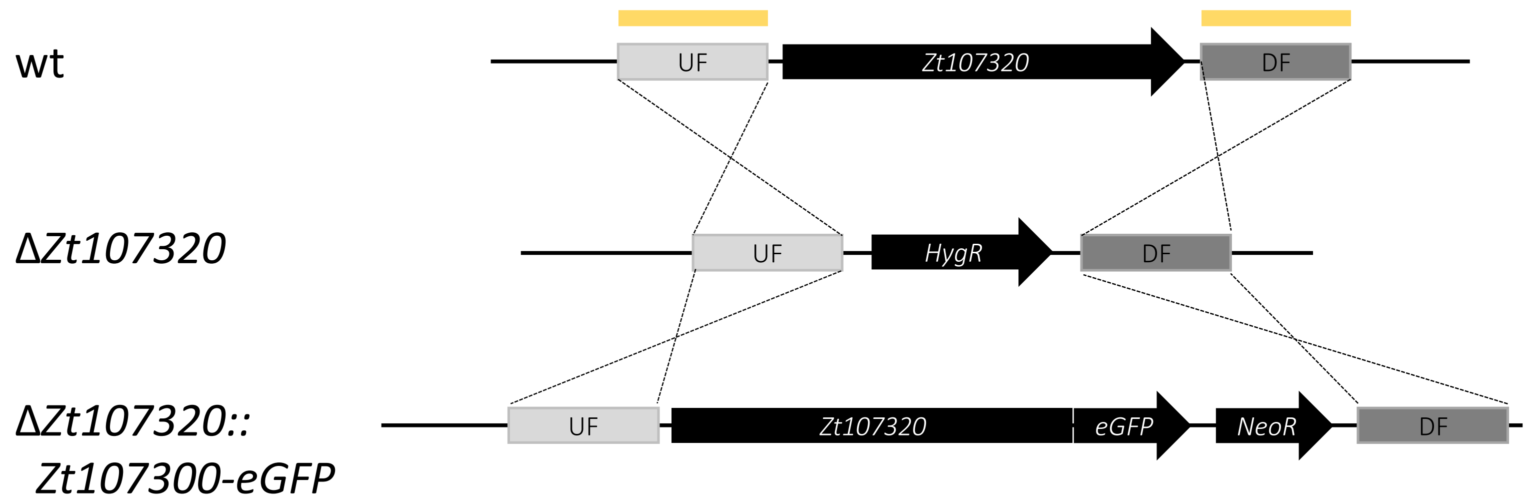


**Figure S3. Generation of *Zt107320* mutants in *Z. tritici***. a) Schematic illustration of the gene replacement strategy of *Zt107320*. Generation of the Δ*Zt107320* strains by homologous recombination between the upstream (UF) and downstream flanking regions (DF) of *Zt107320* in its genomic locus and a plasmid carrying the hygromycin resistance cassette (hygR) located between the UF and DF. Homologous recombination results in the integration of the hygromycin-resistance gene cassette (*hygR*) in the locus of *Zt107320*. Δ*Zt107320::Zt107300-*eGFP were generated by homologous recombination between UF and DF of the Δ*Zt107320* strains and a transformed plasmid containing a c-terminal fusion of *Zt107320* and *eGFP* and a Geneticin resistance cassette (*NeoR*) located between the UF and DF Yellow bars indicate the position of the probes used in the Southern blot analyses.

b)

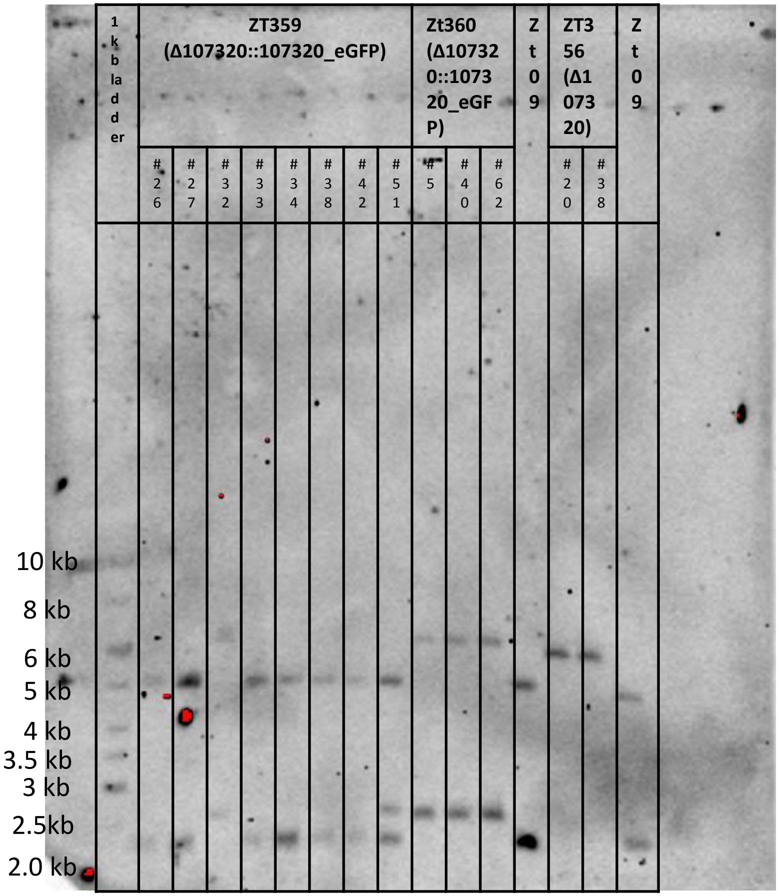


PvuII-Digestion

| Strain | Genotype | Expected bandsizes |
|---|---|---|
| Zt359 | *ΔZt107320::Zt107320_eGFP* | 6237 bp<br>2567 bp |
| Zt360 | *ΔZt107320::Zt107320_eGFP* | 6237 bp<br>2567 bp |
| Zt356 | *ΔZt107320* | 5995 bp |
| Zt09 | wt | 4991 bp<br>2246 bp |

Correct strains:
ZT356#20 = ΔZt107320 strain 1
ZT356#38 = ΔZt107320 strain 2

ZT360#5 = ΔZt107320::Zt107320_egfP strain1 = Zt107320 complement strain 1
ZT360#40 = ΔZt107320::Zt107320_egfP strain2 = Zt107320 complement strain 2

**Figure S3. Generation of *Zt107320* mutants in *Z. tritici*.** b) Confirmation of *Zt107320* mutants by Southern blot analyses.
